# The small GTPase Ran defines Nuclear Pore Complex Asymmetry

**DOI:** 10.1101/2024.10.10.617378

**Authors:** Jenny Sachweh, Mandy Börmel, Sven Klumpe, Anja Becker, Reiya Taniguchi, Marta Anna Kubańska, Verena Pintschovius, Eva Kaindl, Jürgen M. Plitzko, Florian Wilfling, Martin Beck, Bernhard Hampoelz

**Author notes:** Department of Biology, Technical University of Darmstadt, Darmstadt, Germany (Verena Pintschovius).

## Abstract

Nuclear pore complexes (NPCs) bridge across the nuclear envelope and mediate nucleocytoplasmic exchange. They consist of hundreds of nucleoporin building blocks and exemplify the structural complexity of macromolecular assemblies. To ensure transport directionality, different nucleoporin complexes are attached to the cytosolic and nuclear face of the NPC. How those asymmetric structures are faithfully assembled onto the symmetric scaffold architecture that exposes the same interaction surfaces to either side, remained enigmatic. Here we combine cryo-electron tomography, subtomogram averaging, and template matching with live cell imaging to address this question in budding yeast and *Drosophila melanogaster*. We genetically induce ectopic nuclear pores and show that pores outside the nuclear envelope are symmetric. We furthermore demonstrate that the peripheral NPC configuration depends on the nucleotide state of the small GTPase Ran. Our findings indicate that the nuclear transport system is self-regulatory, namely the same molecular mechanism controls both transport and transport channel composition.

## Introduction

Eukaryotic cells use nuclear pore complexes (NPCs) to mediate and regulate nucleocytoplasmic transport across the nuclear envelope (NE). NPCs are ∼60-120 MDa macromolecular complexes composed of multiple copies of about 30 nucleoporins (Nups). These form biochemically stable subcomplexes that assemble into an eightfold symmetric hollow cylindrical structure at fusion sites between the inner and the outer nuclear membrane (Figure 1A). Recent advances in cryo-electron tomography (cryo-ET) and integrative modelling have led to an increasingly complete structural model of the NPC ^1^. The structured scaffold is the anchor point for more flexible regions such as the intrinsically disordered Phenylalanine-Glycine (FG) domains in Nups which enable selective transport through the NPC’s central channel. Directionality of active transport across the NPC is facilitated by the differential nucleotide states of the small GTPase Ran in the cytoplasm and the nucleus ^2^.

**Figure 1:**
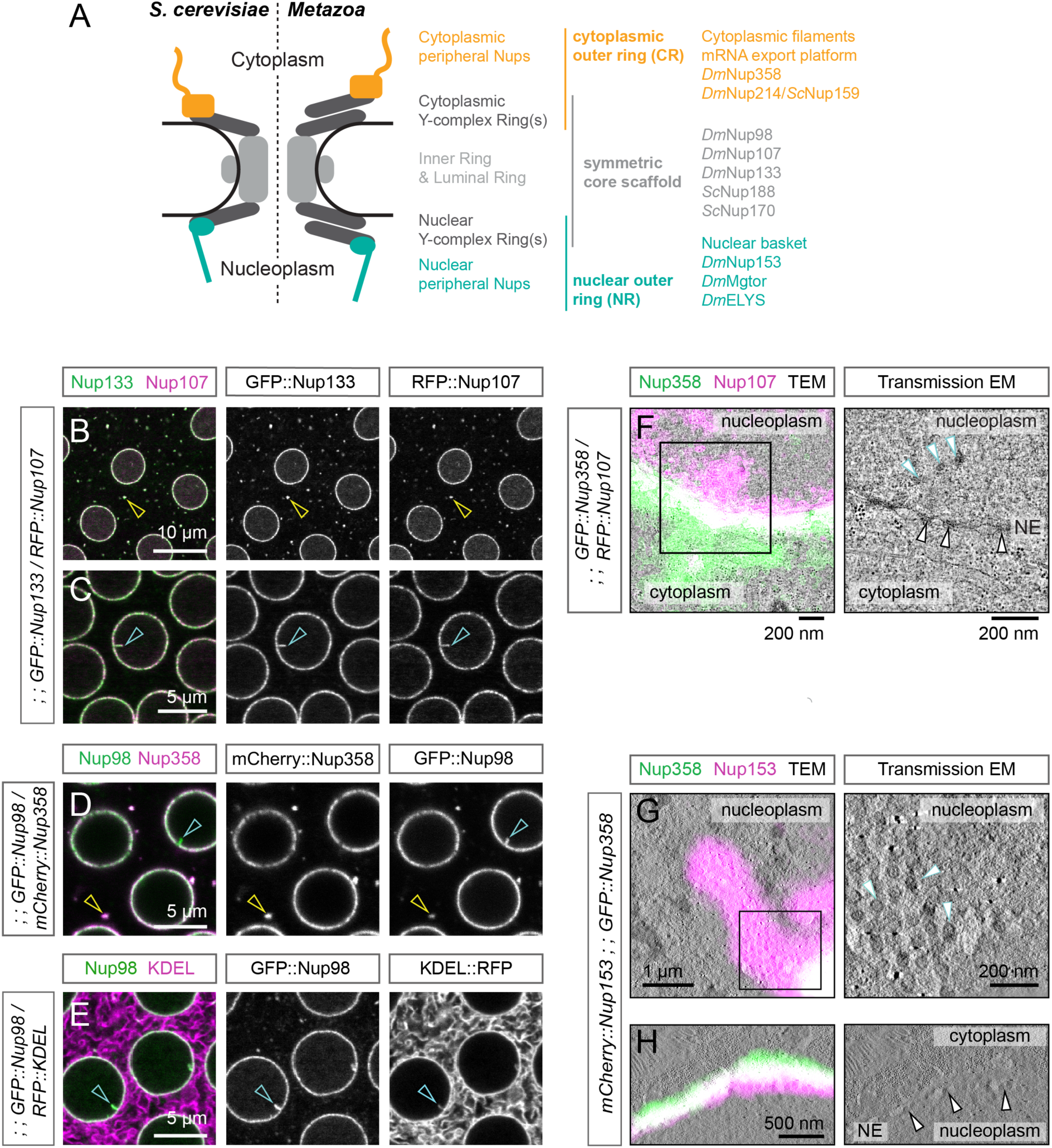
NPC composition depends on the subcellular context. **A** Schematic representation of the structured scaffold of a nuclear pore complex in *S. cerevisiae* and metazoa, highlighting asymmetric structures exclusive to the nuclear or cytoplasmic side on the symmetric core scaffold in teal and orange, respectively. The location of yeast (*Sc*) or *Drosophila* (*Dm*) nucleoporins used in this study and mentioned substructures are indicated. **B-E**: Distinct localization of different nucleoporins. Confocal images of *Drosophila* syncytial blastoderm embryos of the indicated genotype. **B, C** The NPC’s core scaffold components GFP::Nup98 and RFP::Nup107 localize to the nuclear envelope, to cytoplasmic AL (yellow arrowhead in B) and to intranuclear stretches projecting out from the NE (cyan arrowhead in C). **D** The peripheral cytoplasmic mCherry::Nup358 localizes to the NE, cytoplasmic AL (yellow arrowhead), but not to intranuclear GFP::Nup98-positive stretches (cyan arrowhead). **E** Intranuclear Nup98 stretches contain the membrane marker KDEL::RFP (cyan arrowhead). **F** Intranuclear Nup stretches are correspond to nucleoplasmic NPCs. In-section CLEM of the nuclear periphery of a *Drosophila* syncytial blastoderm embryo expressing fluorescently tagged Nups. RFP::Nup107 colocalizes with electron-dense NPCs in the NE (black arrowheads) and in the nucleoplasm (cyan arrowheads). GFP::Nup358 localizes to the cytoplasmic side at the NE but is absent from nNPCs. Right panel shows inset indicated in the overlay image. **G-H** nNPCs contain peripheral nucleoplasmic Nups. In-section CLEM of the periphery of a nurse cell nucleus in an egg chamber dissected from a female expressing GFP::Nup358 and mCherry::Nup153. **G** Top views of densely packed nNPCs (cyan arrowheads) that are positive for mCherry::Nup153 but are devoid of GFP::Nup358. Right panel shows inset indicated in the overlay image. **H** Both tagged proteins are present at NPCs at the NE (arrowheads). See also Figure S1.

The NPC’s structured scaffold lining the pore membrane consists of three stacked rings (Figure 1A): cytoplasmic (CR), inner (IR) and nuclear (NR) ring. NPCs are surrounded by a circular arrangement of a single Nup in the NE lumen that connects to the IR via transmembrane domains ^3^. This luminal ring and the inner ring are symmetric across the NE membrane plane, while different outer rings (NR and CR) attach to the IR on either side of the membrane (Figure 1A). In both outer rings, Y-complexes (also called Nup107-160 complexes) form IR-proximal ring-shaped substructures, which are bound by distinct compartment-specific peripheral Nups (Figure 1A). The cytoplasmic ring contains the evolutionary conserved Nup214/Nup88 (yeast Nup159) subcomplex, which forms the mRNA export platform ^4^ and the metazoan specific Nup358/RanBP2 ^5,6^ (Figure 1A). On the nuclear side, the peripheral Nups ELYS/Mel28, Nup153, TPR, Nup50 and ZC3HC1 ^7–11^ in metazoans and Mlp1, Mlp2, Nup1, Nup2, Nup60, Pml39p and Ely5 in yeast form nuclear-specific structures, such as the nuclear basket ^12,13^ (Figure 1A). The compositional asymmetry of NPCs across the nuclear membrane has been extensively documented and is known to be critical for cellular functions such as mRNA export, directional protein transport or genome organization ^14^. However, it remains unclear how asymmetry is established on the underlying symmetric structure of the core scaffold. It is equally unknown if, in principle, NPCs could exist in a configuration that is symmetric across the NE plane.

To address how this asymmetric configuration of the NPC is regulated, we characterized ectopic NPCs within double membranes outside of the NE, which are known to differ in composition from those in the nuclear envelope. In rapidly dividing cells, cytoplasmic NPCs (cNPCs) populate specialized endoplasmic reticulum (ER) sheets called Annulate Lamellae (AL) (reviewed in ^15^). In turn, nucleoplasmic NPCs (nNPCs) have been reported to occur in membranes within the nucleoplasm ^16^. We resolve the structures of the three varieties, (NE-)NPC, cNPC and nNPC, from two organisms, *Drosophila melanogaster* and *Saccharomyces cerevisiae*, and demonstrate that the NPC core scaffold can be established with a symmetric configuration of peripheral nucleoporins. We further found that the NPC adopts its peripheral configuration according to the subcellular environment, where the small GTPase Ran acts as a key regulator. This suggests that NPC composition dynamically adapts to the surrounding environment, ensuring the correct, asymmetric architecture of NPCs in the nuclear envelope.

## Results

### NPC composition depends on the subcellular context

Annulate Lamellae are abundant in *Drosophila* syncytial blastoderm embryos and insert into the NE during the rapid interphases before cellularisation ^17^. They have also been described to contribute to postmitotic nuclear envelope re-assembly in human cells ^18^. Nups of the NPC core scaffold and the cytoplasmic component Nup358, but not peripheral nuclear Nups, were found at AL ^17,19,20^. However, whether AL-NPCs are symmetric structures remained unknown.

To study NPC composition outside of the NE context, we imaged syncytial blastoderm embryos from transgenic fly lines expressing fluorescently tagged Nups. As expected, the NPC core scaffold components Nup107, Nup133 and Nup98 localize to the NE and to AL (Figure 1B-D). Moreover, all three proteins label stretches projecting from the NE into the nucleoplasm (Figure 1C-D). The cytoplasmic peripheral Nup358 localizes to the NE and AL, but not to intranuclear Nup98-positive stretches (Figure 1D). Colocalization with the fluorescent ER and NE membrane marker RFP::KDEL shows that these stretches contain membranes, indicating that the labelled Nup might be part of nucleoplasmic NPCs (Figure 1E). In-resin correlative light and electron microscopy (CLEM) on high-pressure frozen blastoderm embryos indeed confirms the presence of nNPCs at intranuclear membrane stretches. These nucleoplasmic NPCs colocalize with RFP::Nup107, but are devoid of GFP::Nup358 (Figure 1F). Absence of the mRNA export platform constituent Nup214::GFP from nNPCs further supports our conclusion that cytosolic Nups are absent from nucleoplasmic NPCs (Figure S1A-B).

We next asked if nNPCs in turn contain peripheral Nups specific to the nuclear side of canonical NE-NPCs. We performed CLEM experiments on *Drosophila* ovaries expressing the peripheral nuclear Nup mCherry::Nup153 and the peripheral cytoplasmic Nup GFP::Nup358. nNPCs indeed contain mCherry::Nup153, but not GFP::Nup358 (Figure 1G), while both components are present at the NE (Figure 1H). In embryos, nNPCs contain the nuclear peripheral nucleoporins Nup153 and Mgtor (the fly TPR homolog). Both are absent from AL, similar to the nuclear peripheral nucleoporin ELYS (Figure S1A-C). Thus, we conclude that NPCs in *Drosophila* exhibit a distinct composition depending on their subcellular localization: NE-NPCs contain both sets of peripheral nucleoporins, while cNPCs and nNPCs, which are exposed to the same compartment on both sides, only associate with either cytoplasmic or nuclear peripheral Nups, respectively. However, while this shows that cNPCs and nNPCs contain only the compartment-appropriate set of peripheral Nups, it is not clear whether the resulting NPCs contain peripheral Nups on only one or both sides, and thus constitute symmetric or asymmetric structures.

To quantify the relative abundance of core and peripheral Nups in cNPCs, we compared the mean fluorescence intensity ratios of co-expressed tagged Nups between AL and NE (Figure S1D-E). The ratios of the core components RFP::Nup107, GFP::Nup98 and GFP::Nup133 at the NE and AL are similar, while relative to these core components, the cytoplasmic peripheral nucleoporin mCherry::Nup358 is more abundant at AL compared to the NE (Figure S1D-E). At the same time, the ratio of mCherry::Nup358 to Nup214::GFP, another cytoplasmic peripheral nucleoporin, is similar at AL and at the NE (Figure S1E). These data suggest that Nup358 and Nup214 are more abundant at cNPCs than at NE-NPCs, which would be consistent with a model of symmetric cNPCs that exhibit two cytoplasmic rings.

### Enforced dimerization of Nup358 induces Annulate Lamellae

To directly validate a symmetric configuration of peripheral Nups on cNPCs, as suggested above we sought to enrich AL for structural analysis. In the female *Drosophila* germline (Figure S2A), condensation of Nup358 in nurse cells is a prerequisite to assemble cNPCs at AL in the ooplasm. Depletion of *nup358* interferes with AL assembly and integrity ^21^. We thus reasoned that enforcing additional interactions between Nup358 molecules could enhance AL formation. *Drosophila* Nup358 domain architecture resembles that of human Nup358 ^22^, which is anchored to the cytoplasmic outer ring via its folded N-terminal domain ^23–25^. The remaining part of the protein, which is mainly composed of alternating Ran-binding domains and intrinsically disordered regions, extends into the cytoplasm ^24,26,27^. We generated and visualized Nup358-Nup358 dimers by endogenously fusing one of the two parts of the split-Venus fluorescent protein, Venus N-terminus (VN) and C-terminus (VC), to the Nup358 C-termini ^28^. In offspring of crosses between the two lines, quasi-irreversible bond formation between VN and VC produces the fluorescent Venus protein, and dimerizes Nup358. Male and female flies that co-express both transgenes are viable, but sterile.

In egg chambers from Nup358::VN/Nup358::VC females, Venus fluorescence is detected at the NE and at ooplasmic AL, as expected (Figure 2A). However, it is also present in large non-spherical clusters indicative of AL in the cytoplasm of nurse cells (Figure 2A). This differs from the typical spherical Nup358 condensates found in nurse cells of non-dimerizing Nup358::GFP egg chambers or of wild type egg chambers stained for *Dm*Nup358 (Figure 2B-C). CLEM experiments confirm that the Venus-positive aggregates in the cytoplasm of Nup358::VN/Nup358::VC nurse cells correspond to AL populated with cNPCs (Figure 2D).

**Figure 2:**
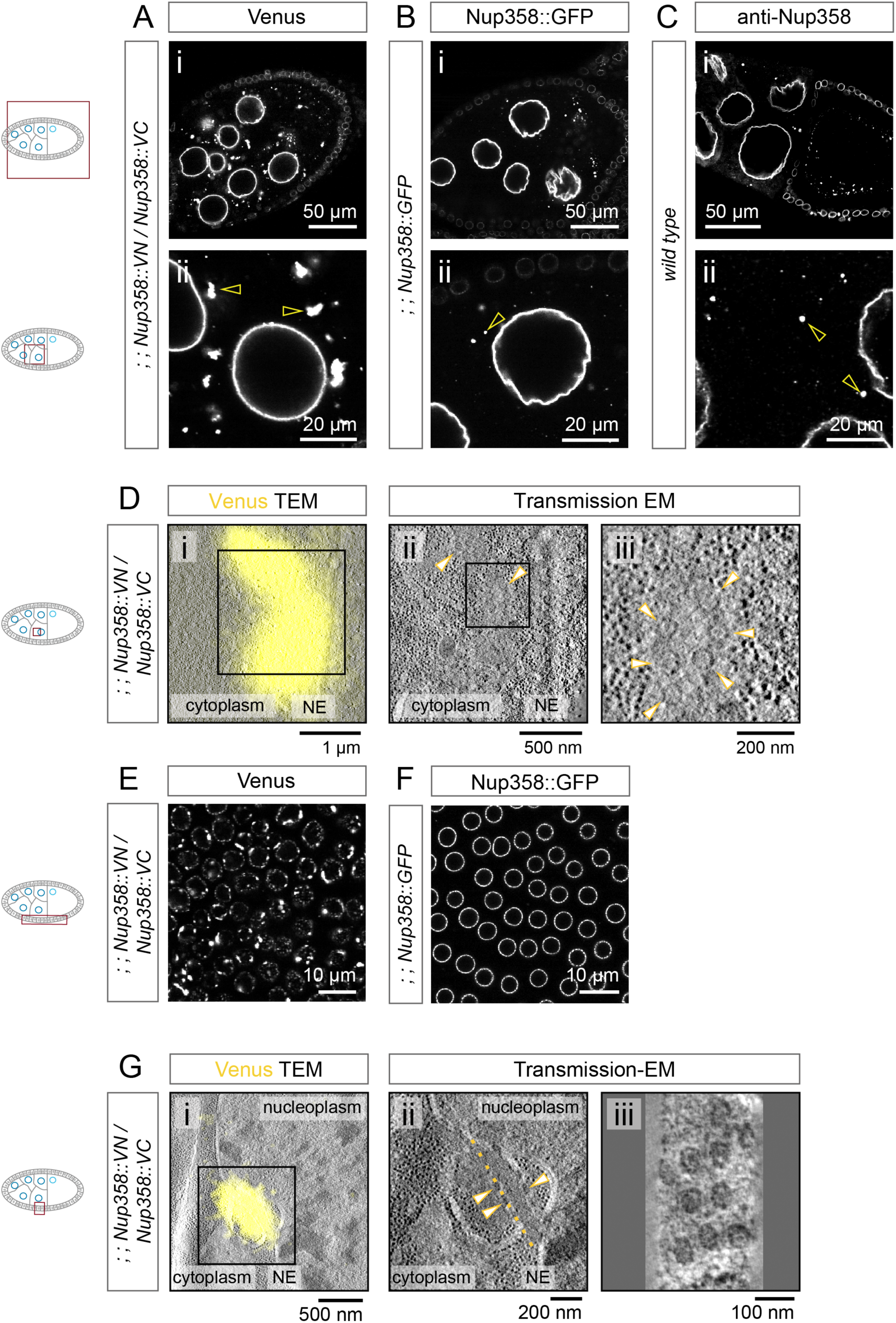
Enforced dimerization of Nup358 induces Annulate Lamellae. A-C Enforced dimerization of Nup358 induces cytoplasmic Nup358 foci in nurse cells. Confocal images of *Drosophila* egg chambers (i) or nurse cells (ii) from ovaries of Nup358::VN/Nup358::VC (A), Nup358::GFP (B) or wild type (C) females. **A** Large Venus fluorescent foci are present in the cytoplasm of Nup358::VN/Nup358::VC ovaries (arrowheads in A ii). **B-C** In Nup358::GFP control ovaries, fluorescent signal is present at the NE and in small spherical condensates (arrowheads in B ii), similar to wild type ovaries immunostained for Nup358 (arrowheads in C ii). **D** Venus foci in Nup358::VN/ Nup358::VC nurse cells are Annulate Lamellae. In-section CLEM of a *Drosophila* Nup358::VN/Nup358::VC egg chamber. Venus fluorescence (i) correlates with a ribosome-free cytoplasmic patch populated with cNPCs (yellow arrowheads in ii, iii). Slicing through the tomographic volume at a different angle highlights cNPCs in top views (iii). (ii) corresponds to the boxed region in (i), (iii) to the boxed region in (ii). **E, F** Enforced Nup358 dimerization induces cytoplasmic Nup358 foci in follicle cells. Confocal images of follicle cells in dissected ovaries of either Nup358::VN/Nup358::VC (E) or Nup358::GFP control flies (F). Fluorescent cytoplasmic or NE-attached foci accumulate upon Nup358 dimerization (E) but not in Nup358::GFP follicle cells, where GFP is exclusively at the NE (F). **G** Venus foci in Nup358::VN/Nup358::VC follicle cells are Annulate Lamellae. In-section CLEM of a Nup358::VN/Nup358::VC egg chamber. Venus signal correlates with an area containing AL adjacent to the nucleus of a follicle cell (i). Tilted tomographic slice (ii) corresponding to the boxed region in (i) indicates cNPCs in membranes (arrowheads). (iii) Magnified tomographic slice perpendicular to the normal of the membrane indicated in yellow in (ii) highlights cNPCs in top views. See also Figure S2.

In wild type ovaries, nucleoporin condensation and AL formation is prominent in the germline but largely absent from the surrounding somatic follicle cells ^21^. However, Nup358::VN/Nup358::VC females exhibit intense fluorescent foci at the NE and in the cytoplasm of follicle cells, different from control egg chambers (Figure 2E-F). Those foci correlate with cNPCs in CLEM experiments (Figure 2G). We conclude that Nup358 dimerization promotes ectopic AL formation in both germline and somatic cells.

### Insect and vertebrate NPCs share major structural features

To structurally analyse NPCs at different subcellular localizations, we dissociated Nup358::VN/Nup358::VC ovaries, enriched follicle cells by filtration and prepared them for *in situ* cryo-ET by focused-ion-beam (FIB) milling, as previously described ^29^. Tilt series were acquired around the NE, and both NE-NPCs and cNPCs were readily identifiable in the tomographic reconstructions (Figure 3A).

**Figure 3:**
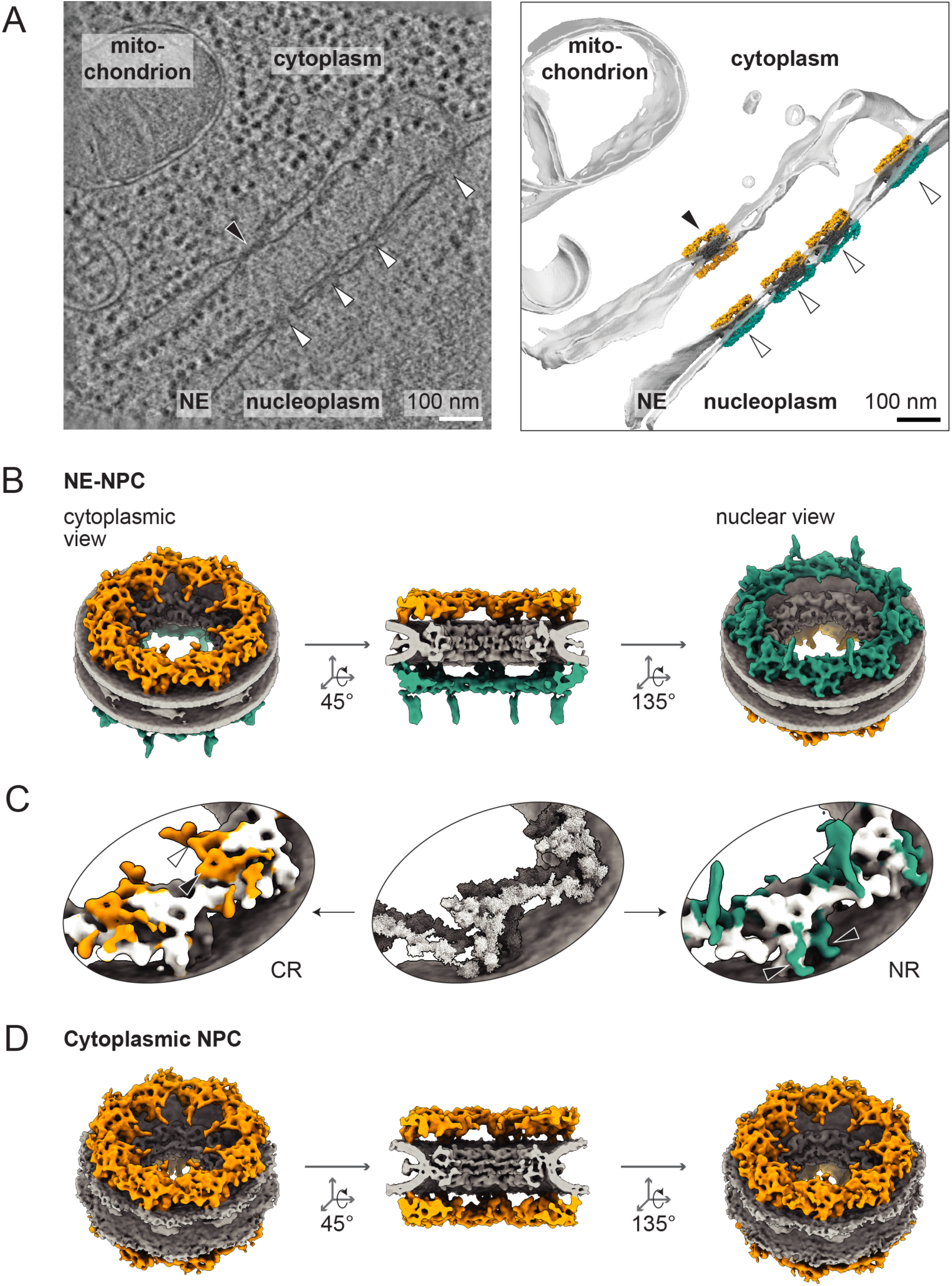
The *in situ* structure of *Drosophila* NPCs resembles vertebrate NPCs and depends on the cellular environment. **A** *In situ* cryo-ET reveals NPCs at the NE and at cytosolic membranes. Slice of a tomogram (left) and membrane segmentation with subtomogram averages mapped back to their locations of origin (right). A cNPC (black arrowhead) is embedded in a cytosolic membrane sheet that is continuous with the NE. NE-NPCs (white arrowheads) decorate the NE. **B** *Drosophila* NE-NPC shown as an 8-fold symmetric composite map in cytoplasmic (left), side-cut (centre) and nuclear view (right). The membrane, inner ring and luminal ring are shown in light grey, the cytoplasmic ring in orange, the nuclear ring in teal. **C** Asymmetry of the *Drosophila* NE-NPC. Zoom-ins of CR (left) and NR (right) of the NE-NPC are shown together with the human double Y-complex arrangement (centre, PDB 7R5J) that fits the *Drosophila* outer ring average densities. Surfaces of the CR and NR averages are coloured according to the fitted double Y-complexes when within a distance of 15 Å to the models. Remaining orange and teal surfaces cannot be explained by Y-complex structures and partially correspond to known binding sites of peripheral Nups on the respective outer rings. The CR (left panel) exhibits densities where the mRNA export platform (white arrowhead) and the base of the cytoplasmic filaments (black arrowhead) would be expected. The NR (right panel) exhibits densities where the nuclear basket (white arrowhead) and the ELYS binding sites (black arrowheads) would be expected. **D** cNPCs have two cytoplasmic outer rings. *Drosophila* cNPC shown as an 8-fold symmetric composite map in top view (left), side-cut view (centre) and bottom view (right). The membrane, inner ring and luminal ring are shown in light grey, the two cytoplasmic rings in orange. **A-D:** Sample: Cells isolated from Nup358::VN/Nup358::VC *Drosophila* ovaries. Colours: CR - orange, NR – teal, IR - grey. See also Figure S3.

We first focused on NE-NPCs in order to assess the structure of an insect NPC. The overall NE-NPC architecture as revealed by subtomogram averaging in *Drosophila* cells closely resembles that of human NPCs ^30^: the inner ring is adjoined by two outer rings, each built from two layers of Y-complexes (Figure 3B, Figure S3A), similar to all metazoan NPCs described to date ^1,31^. At the plane of the inner ring, densities in the lumen of the NE indicate the presence of a luminal ring as known from vertebrate NPC structures ^30^. A CR-specific density extends towards the central channel and is reminiscent in shape and location of the mRNA export platform reported in other species ^32,33^. The densities on top of the double ring of Y-complexes likely correspond to the base of cytoplasmic filaments formed by Nup358 in humans (Figure 3C) ^23,24,30^ or *Xenopus laevis* ^25^. On the nuclear ring, an elongated density pointing towards the nucleoplasm indicates the presence of the nuclear basket. Furthermore, a defined density is apparent where ELYS is known to bind to the nuclear ring in vertebrates ^24,30^. Both of these densities are absent from the cytoplasmic ring (Figure 3C). These data show that insect NPCs are structurally similar to vertebrate NPCs, including the asymmetrically associated nucleoporin subcomplexes. This insect NPC architecture is consistent with the distribution of core and peripheral nucleoporins observed by light microscopy (Figure 1, Figure S1).

### Cytoplasmic NPCs contain two cytoplasmic rings

We next determined the structure of NPCs entirely surrounded by cytoplasm. cNPCs in the Nup358::VN/Nup358::VC cells are embedded in cytosolic membrane sheets. Those are continuous with the outer nuclear membrane and decorated with ribosomes indicating that these membranes are part of the ER (Figure 3 A). The core scaffold architecture of the cNPCs is similar to that of NE-NPCs, including an inner, a luminal, and two outer rings (Figure 3D, Figure S3A), albeit in a slightly more constricted state (Figure S3B-C). Importantly, however, both outer rings of the cNPC average recapitulate the cytoplasmic ring architecture of the canonical NE-NPC (Figure 3D). They exhibit densities indicating the presence of the mRNA export platform and of the cytoplasmic filaments. Neither the nucleus-specific ELYS nor the nuclear basket were present in outer rings of the cNPC (Figure 3C-D). Immunofluorescence experiments furthermore corroborate that ELYS is absent from Venus-positive AL in Nup358::VN/Nup358::VC germline and somatic cells, while localizing to the NE as expected (Figure S2B-C).

In summary, our data indicate that cNPCs are symmetric and are comprised of an IR and two CRs. However, as the averaging approach required initial manual assignment of NPC orientation, we cannot formally exclude that single NPCs are asymmetric, e.g. carrying only one ring with the full set of peripheral Nups. In this scenario, inconsistent assignment of initial NPC orientations might result in a fallaciously symmetric average.

To assess the structure of cNPCs at the level of single outer rings, we performed template matching, an analysis based on constrained cross correlation (CCC) which detects structural features of the provided templates in tomographic volumes independently of averaging ^31,34^. We used the average maps of the NE-NPC’s nuclear and cytoplasmic ring asymmetric unit as search templates to identify positions and orientations in the tomograms that match the template densities (Figure 4A and Video S1).

**Figure 4:**
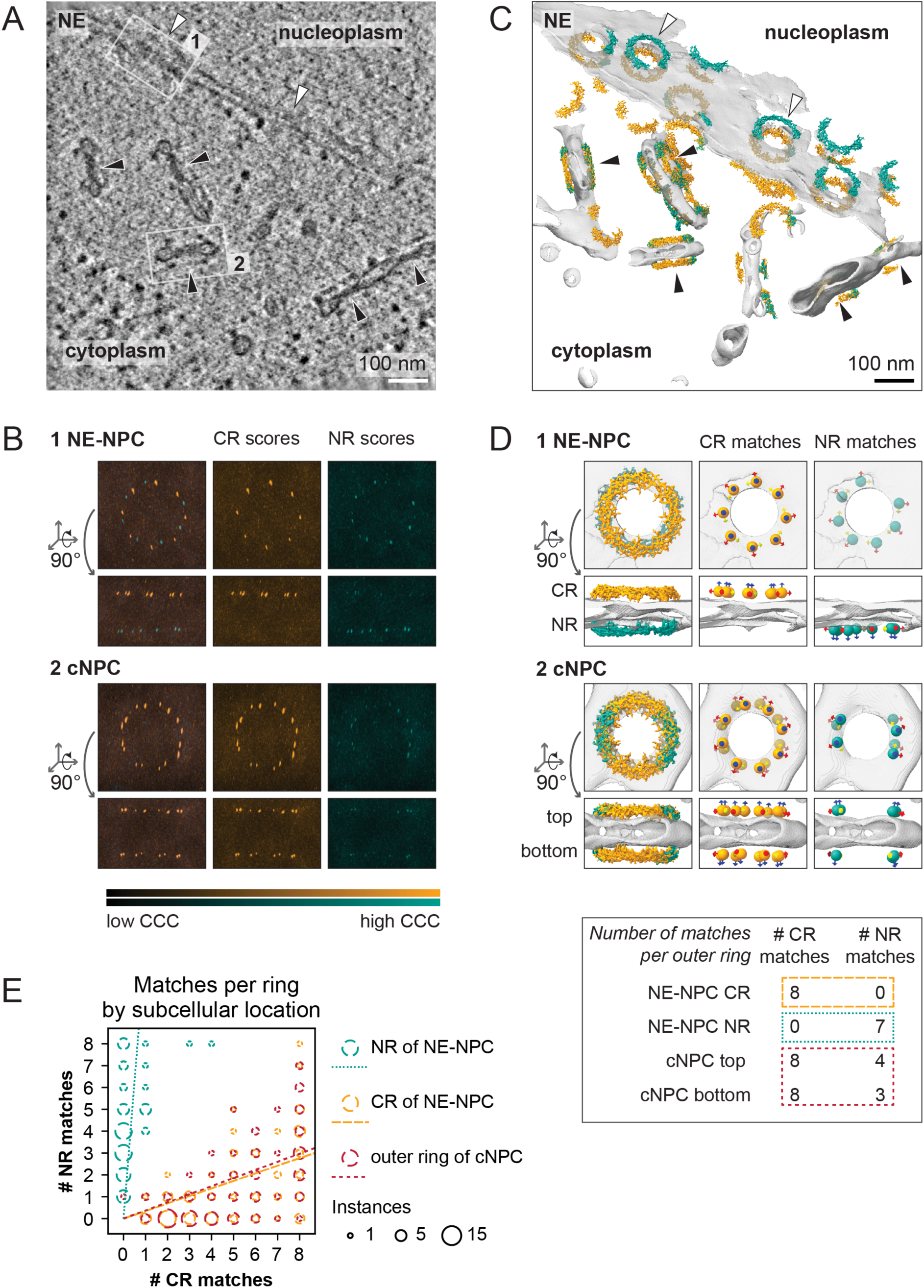
Template matching confirms that cNPCs contain two cytoplasmic rings. A Accumulation of cNPCs close to the NE. Slice from a tomogram showing cNPCs (black arrowheads) and NE-NPCs (white arrowheads). White numbered rectangles designate NPCs presented in subsequent panels. **B** Template matching reveals differences in outer ring symmetry between NE-NPCs and cNPCs on the single NPC level. CCC score volumes obtained from an NE-NPC (location 1 in A) and a cNPC (location 2 in A) are shown in top and side views as maximum-intensity projections. Peaks in the CCC scores indicate a high cross-correlation with the respective template. The NE-NPC has two different outer rings, a nuclear and a cytoplasmic ring which show distinct scores with the two templates, whereas the cNPC features two cytoplasmic rings. **C** Matches of the two templates are enriched in the respective compartments. Membrane segmentation of the same tomogram as (A) with the CR and NR templates mapped back to locations of CCC score peaks. Black and white arrowheads indicate the same NPCs as in (A). **D** Counting matches per outer ring enables quantitative comparisons. Mapped back template matches are shown for the subvolumes presented in (B). Matches are either indicated by mapped back templates or spherical markers. In the table, the number of matches obtained with the two templates is shown per outer ring displayed above. Coloured boxes indicate which group the respective outer ring is assigned to in (E). **E** cNPC outer rings show the same behaviour in template matching as CRs of NE-NPCs. Bubble chart of the matches obtained with the NR template and the CR template per outer ring, grouped by subcellular location. Dashed lines indicate the proportionality of NR vs. CR template matches for each subcellular location. **A-E**: Sample: Cells isolated from Nup358::VN/Nup358::VC *Drosophila* ovaries. Colours: CR - orange, NR - teal. Averages of an asymmetric units of the NE-NPC’s CR and NR (Figure 3B-C) were used as search templates. See also Figure S4 and Video S1.

In a subvolume containing an individual NE-NPC, template matching with the nuclear and cytoplasmic ring templates yielded one eightfold-symmetric ring of high CCC scores in the respectively expected compartment (Figure 4B, panel 1). Weaker CCC peaks detected on the opposite sides are likely due to common structural features of CR and NR. Biological heterogeneity and technical limitations lead to unequal detection of the eight asymmetric units of an outer ring by both templates. Nonetheless, the observed asymmetry of the CCC score peaks across the NE in an NE-NPC subvolume shows that our approach can differentiate nuclear from cytoplasmic outer rings.

Template matching with the same CR template in a cNPC-containing subvolume yields two equally pronounced rings of peaks, indicating that this individual cNPC is symmetric with two cytoplasmic rings (Figure 4B). Consistently, matching with the NR template only results in weak peaks (Figure 4B, panel 2).

By projecting the best matching positions and orientations for the two templates back into the tomographic volume, we can visually assess their distribution (Figure 4C-D). Counting the number of subunit matches with both templates allows to quantitatively compare outer rings grouped by their subcellular location (Figure 4D-E). For the NE-NPC’s NR, the NR template consistently detects more subunits than the CR template, and vice versa. The number of subunits detected by the NR template versus the CR template per outer ring of a cNPC closely recapitulates the observations for the CR of NE-NPCs, again arguing for the presence of two CRs.

Taken together, subtomogram averages and template matching results show that NPCs fully surrounded by cytoplasm are symmetric across the membrane plane, with both outer rings resembling the canonical NE-NPC’s cytoplasmic ring.

### Nucleoplasmic NPCs contain two nuclear rings

Having detected symmetric NPCs in the cytoplasm, we next wanted to test if this is compartment-specific. We used template matching to assess the few nucleoplasmic NPCs in our *Drosophila* data set (Figure S4A-C). Most of the outer rings of nNPCs exhibited more NR than CR template matches on either side of the double membrane, suggesting that nNPCs have two nuclear rings (Figure S4A-C). To exclude that this is a fly specific phenomenon, we next analysed nucleoplasmic NPCs in *S. cerevisiae*. Here, an enrichment of NPCs at intranuclear membrane deformations has been described for multiple mutant strains, including several nucleoporin knockouts ^35^. However, as nucleoporin perturbation likely affects NPC architecture, we instead used a strain lacking *NOT4*. In this background, fluorescently tagged Nup188 and Nup170 form foci, which correspond to clustered NPCs in the NE and in aberrant intranuclear double membranes (Figure 5A-B).

**Figure 5:**
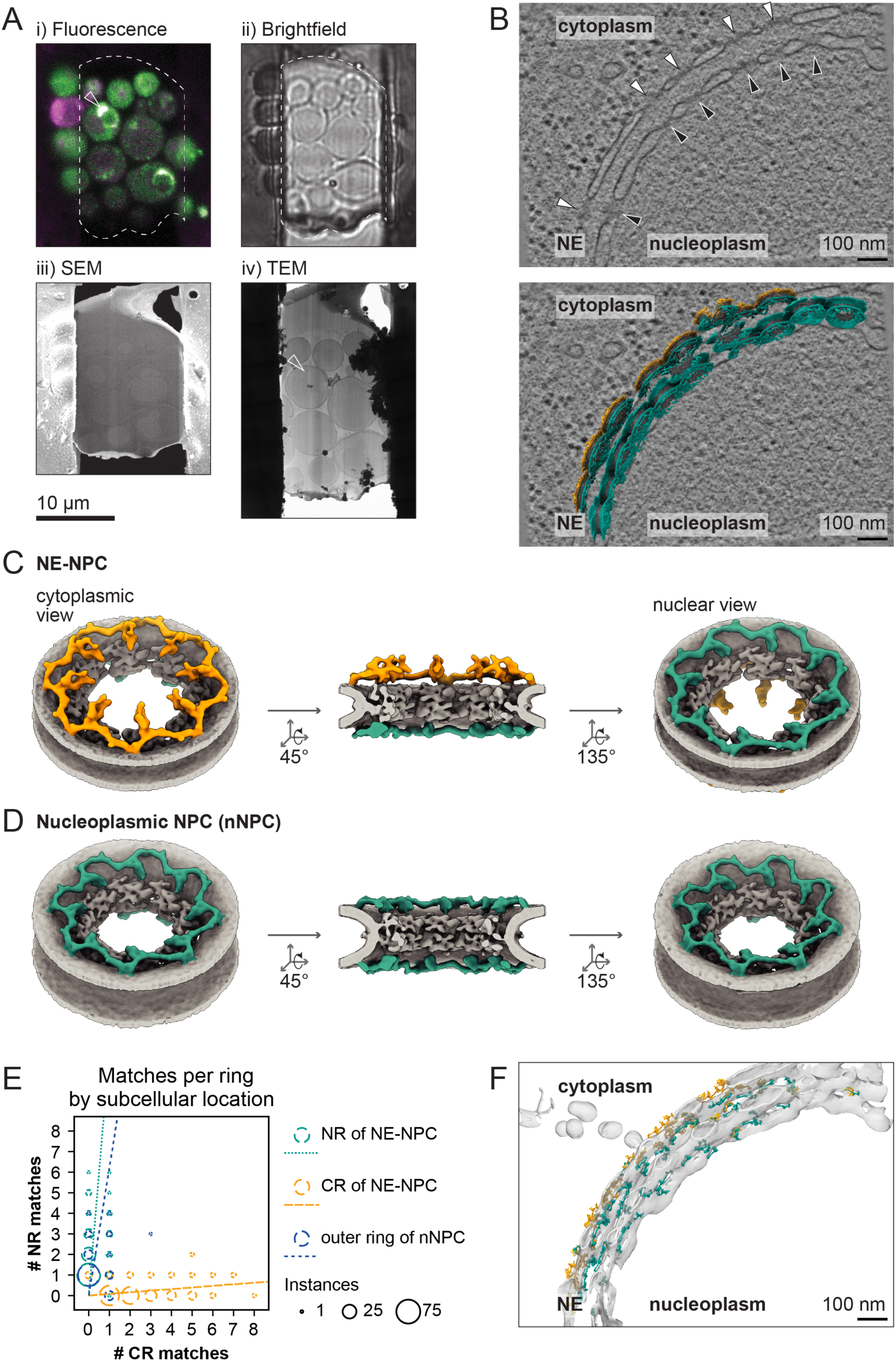
Nucleoplasmic NPCs contain two nuclear rings. **A** Identification of fluorescent NPC foci by cryo-CLEM. Light microscopy of rough milled lamellae (i - green: GFP channel, magenta: mars channel, ii - brightfield) allows to selectively thin lamellae of interest by FIB-SEM (iii) and target the tilt series acquisition in TEM (iv). Arrowhead points to fluorescent focus that was targeted in tomogram acquisition **B** Tomogram of the fluorescent focus from (A) contains clustered NPCs: nNPCs (black arrowheads) embedded in an intranuclear membrane sheet and NE-NPCs in the NE (white arrowheads) are visible. Slice from a tomogram (top panel) or subtomogram averages mapped back to their locations of origin (bottom panel) are shown. Only particles in front of the tomographic slice are visible. **C** *S. cerevisiae* NE-NPC shown as an 8-fold symmetric composite map in cytosolic view (left), in side-cut view (centre) and nuclear view (right). The membrane, inner ring and luminal ring are shown in light grey, the cytoplasmic ring in orange, the nuclear ring in teal. The nuclear ring lacks the inward pointing densities present at the cytoplasmic ring that correspond to the mRNA export platform. **D** nNPCs have two nuclear outer rings. *S. cerevisiae* nucleoplasmic NPC shown as an 8-fold symmetric composite map in top view (left), side-cut view (centre) and bottom view (right). The membrane, inner ring and luminal ring are shown in light grey, the two outer rings, which resemble the nuclear ring from NE-NPCs, in teal. **E** nNPC outer rings show the same behaviour in template matching as NRs of NE-NPCs. Bubble chart of the matches obtained with the NR template and the CR template per outer ring, grouped by subcellular location. Dashed lines indicate the proportionality of NR vs. CR template matches for each subcellular location. **F** Membrane segmentation with the CR (orange) and NR (teal) templates mapped back to locations of high CCC scores from the tomogram shown in (B). **A-F** Sample: *S. cerevisiae* Not4Δ Nup188::GFP Nup170::mars. Colours: CR - orange, NR – teal, IR - grey. See also Figure S5.

In line with the architecture of wildtype NE-NPCs *in situ* ^36^, the subtomogram average of NE-NPCs in the *not4* null strain exhibits an inner ring complemented by an outer ring on either side of the membrane, each composed of one layer of Y-complexes, as well as a luminal ring lining the pore membrane (Figure 5C, Figure S5A). Budding yeast lacks orthologues of both *nup358* and *elys*, and no density corresponding to the nuclear basket is visible in our data.

However, the cytoplasm-specific mRNA export platform clearly distinguishes the CR from the NR (Figure 5C). Overall, our structure indicates that *NOT4* deletion does not dramatically perturb NPC architecture. nNPCs exhibit an intact core scaffold similar to the NE-NPCs, consisting of an inner ring, a luminal ring, and two outer rings, both of which are composed of one layer of Y-complexes (Figure 5D). Thus, the nNPC’s composition differs from incomplete NPCs found at the neck of NE herniations, but similar to those is more constricted than NE-NPCs ^36^ (Figure S5B-C). Strikingly, the mRNA export platform density is absent from both outer rings of the nNPC average, suggesting symmetry across the membrane plane with two nuclear rings (Figure 5D). Template matching with the NE-NPC’s CR and NR templates confirms the presence of two nuclear rings in nNPCs (Figure 5E-F).

Our findings demonstrate that symmetric NPCs exist across evolutionary distant species and across subcellular compartments. We thus conclude that the NPC core is able to accommodate both symmetric and asymmetric binding of peripheral nucleoporins, and that peripheral NPC composition is regulated by the surrounding cellular environment, rather than being inherent to the structure.

### The nucleotide state of Ran determines NPC (a)symmetry

To address potential mechanisms that could regulate NPC symmetry, we tested if the small GTPase Ran is involved. Like other proteins, soluble nucleoporins can be transported through the NPC. This relies on Nuclear Transport Receptor (NTR) binding and a gradient of Ran nucleotide states, which drives active transport across the NE. RanGTP is generated in the nucleoplasm ^37^, and import complexes formed in the cytoplasm dissociate upon RanGTP binding in the nucleus ^38^. On the other hand, export complexes formed in the nucleus in the presence of RanGTP disassemble in the cytoplasm upon GTP hydrolysis ^2^. Thus, NTRs bind and chaperone subsets of nucleoporins in a compartment-specific manner, which makes them selectively available for binding to NPCs ^39,40^. This suggests a simple, yet elegant, model, where transport and composition of the transport channel itself are regulated by the same molecular mechanism. This model predicts that interfering with the Ran nucleotide state shall affect the distribution of peripheral Nups at NPCs.

To test this prediction, we analysed the peripheral nucleoporin distribution upon Ran gradient perturbation. We injected the non-hydrolysable GTP analogue GTPγS into *Drosophila* syncytial blastoderm embryos, where due to the absence of cell boundaries injected compounds diffuse freely, resulting in a concentration gradient within the embryo (Figure 6A i). Co-injection of fluorophore-conjugated Wheat Germ Agglutinin (WGA) marks the injection site in the embryo and visualizes NPCs via FG-Nup binding. Strikingly, close to the injection site, the peripheral nuclear Nup Mgtor::mCherry accumulated at GFP::Nup358-positive cytoplasmic AL (Figure 6A ii). In contrast, AL were devoid of Mgtor::mCherry distant from the injection site (Figure 6A iii).

**Figure 6:**
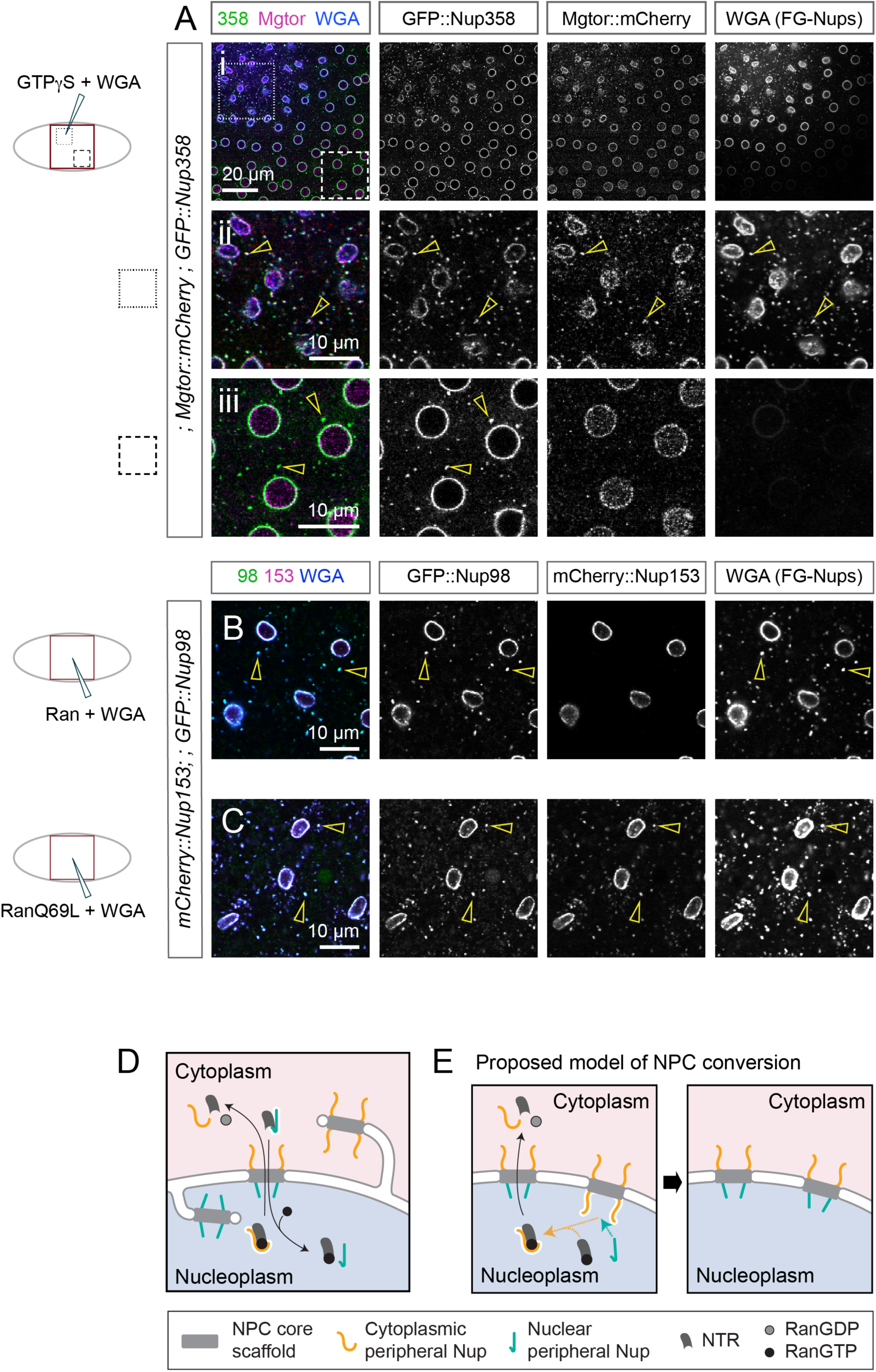
The nucleotide state of Ran determines NPC asymmetry. **A-C** Cytoplasmic RanGTP provokes the localization of peripheral nuclear Nups to AL. Confocal images through the nuclei of *Drosophila* syncytial blastoderm embryos of the indicated genotypes after co-injection of an effector with fluorescently labelled WGA that marks the injection site and visualizes NPCs via FG-Nup binding. Note that there are no cellular borders within the syncytium and injected components diffuse gradually away from the injection site. **A** After GTPγS injection, the nuclear basket component Mgtor::mCherry colocalizes with GFP::Nup358-positive AL near the injection site (ii, arrowheads) but not more distant from the injection site (iii). **B, C** Cytoplasmic GFP::Nup98 stained AL (arrowheads) are devoid of the peripheral nuclear mCherry::Nup153 after injection of recombinant *Hs*Ran (B) but accumulate mCherry::Nup153 upon injection of the hydrolysis-deficient mutant *Hs*RanQ69L (C). **D** A model for the regulation of peripheral NPC composition. The cellular compartment determines peripheral Nup configuration on the NPC’s core scaffold, resulting in asymmetric NE-NPCs and symmetric nNPCs and cNPCs. Ran-NTR mediated chaperoning and transport of peripheral nucleoporins differentiate their availability in the cytoplasm and the nucleoplasm, resulting from selective binding or dissociation of the respective import or export NTRs due to high levels of RanGTP in the nucleoplasm or GTP hydrolysis in the cytoplasm. NTR-bound Nups do not attach to the NPC. **E** Proposed model for NPC conversion after AL insertion. Insertion of cNPCs into the NE exposes a previously cytoplasmic outer ring to the nuclear RanGTP environment. This leads to a compartment-appropriate reconfiguration of the peripheral NPC components.

To directly test if the peripheral Nup composition depends on the nucleotide state of Ran, we next injected either recombinant wild-type *Hs*Ran or hydrolysis-deficient *Hs*RanQ69L ^41^ (Figure 6B-C). Unlike wild-type Ran, injection of the hydrolysis-deficient RanQ69L mutant caused an enrichment of the nuclear peripheral nucleoporin mCherry::Nup153 at AL in the cytoplasm (Figure 6B-C). This corroborates that the Ran nucleotide state determines which set of peripheral Nups associates with the symmetric NPC core.

## Discussion

Functionality of the nuclear pore complex at the nuclear envelope critically depends on its asymmetry across the NE plane ^14,21^. A distinct configuration of nuclear and cytoplasmic peripheral Nups on the symmetric NPC core is for example essential for mRNA export, directional shuttling of protein cargoes across the NE, as well as nuclear transport-independent functions including peripheral chromatin organization. However, how a cell ensures the correct configuration on each side of the nuclear envelope despite the symmetric NPC core scaffold, remained unclear. By analysing NPCs in as well as outside of the NE through orthogonal imaging techniques, we establish that the NPC’s core scaffold *per se* can accommodate binding of peripheral Nups either in an asymmetric or symmetric configuration. We find that the respective configuration is determined by the subcellular compartment and molecularly regulated by the nucleotide state of the small GTPase Ran.

### Context-dependent NPC architecture

Across eukaryotes, nuclear pore complexes share a common eight-fold rotationally symmetric, three-tiered architecture (Figure 1A), yet compositional differences exist not only between species, but also within one organism. The repertoire of nucleoporins can vary between cell types or developmental stages ^42^ as well as between distinct nuclei within the same binucleated cell ^43^. Compositional variation is also possible within the same nucleus: in budding yeast, NPCs adjacent to the nucleolus are devoid of basket components ^44,45^. All these variations concern NPCs in their canonical location on the nuclear envelope. In contrast, NPCs found at the neck of herniations in yeast do not share this configuration suggesting an interdependence of NPC architecture and its environment ^36^.

This raises the question whether the architecture of ectopic NPCs outside the NE would be identical to pores that bridge the nuclear membranes. Here we applied *in situ* cryo-ET to study the structures of NPCs in different cellular compartments in both budding yeast and *Drosophila.* This yielded a detailed map of the so far undescribed insect NPC, which overall resembles that of other metazoan species ^1,31^(Figure 3B), as confirmed by successful template matching with a human search template (Video S1). In both yeast and *Drosophila,* NPCs outside the NE exhibit the same core scaffold architecture as on the NE, including the interaction with the luminal ring across the pore membrane. This highlights that most nucleoporin interactions within the NPC core scaffold do not require localization on an intact NE.

Despite their common core scaffold, NPCs outside the nuclear envelope exhibit a smaller diameter than NE-NPCs (Figure S3B-C and Figure S5B-C). Constitutive tension on the NE is thought to stretch NPCs, while they constrict when tension is relieved ^46–48^. It is conceivable that cNPCs and nNPCs do not experience identical forces to NPCs on the NE, possibly explaining the more constricted state of these pores.

Unique densities characteristic for peripheral Nups of the two outer rings allow us to address if NPC asymmetry is preserved outside the NE. Our *in situ* analysis reveals that NPCs have a symmetric configuration of peripheral Nups when they are embedded in membranes that are surrounded by the same cellular milieu.

### Regulation of NPC (a)symmetry

All NPC structures reported to date are asymmetric across the membrane plane. This asymmetry manifests in different sets of peripheral Nups binding to the core scaffold. In addition, differences in the copy number or composition of Y-complexes ^45,46,48,49^ or in the curvature of the pore membrane ^50^ may distinguish the nuclear and cytoplasmic side ^51^.

In both model systems used in this study the two outer rings are each composed of the same number of Y-complexes, resulting in highly similar interfaces available for peripheral Nup binding on either side of the NPC. To nevertheless yield an asymmetric peripheral Nup configuration, NPC-intrinsic or extrinsic mechanisms would be plausible. An NPC-intrinsic model would require structural determinants that result in different binding site availabilities for peripheral Nups. Indeed, steric occlusion prevents simultaneous binding of some, but not all, peripheral Nups in the case of the human NPC ^24^. To explain asymmetry across the NE, peripheral Nup configuration on one outer ring would have to invoke a modification of the respective interaction sites on the opposing outer ring. According to such a model, binding of the same peripheral Nups on both sides of an NPC would be prevented.

However, our data demonstrate that symmetric NPCs with the same peripheral components exist in cells, strongly arguing against an NPC-intrinsic model. Instead, our results speak for an alternative mechanism, where the NPC core is a versatile element that adopts the “correct” set of peripheral Nups in response to the cellular compartment it is exposed to. In our experiments elevated levels of cytoplasmic RanGTP caused an alteration of peripheral Nup composition on cNPCs (Figure 6A-C), suggesting that the nucleotide state of the small GTPase Ran is a critical determinant of the NPC’s configuration. We thus conclude that extrinsic cues regulate NPC asymmetry.

This effect could be mediated by the nucleocytoplasmic transport system itself: active nucleocytoplasmic transport depends on the RanGTP gradient across the NE. Nucleoporins are recognized by NTRs, which transport them across the NE, leading to compartment-specific enrichment ^52,53^, which is crucial for interphase NPC assembly ^54,55^. In addition, binding to NTRs limits the propensity of Nups to condense or aggregate ^56^, a mechanism that is also employed when the NPC is dismantled during open mitosis ^40^. Consistent with this, NTRs negatively regulate NPC assembly, indicating that only Nups that are not chaperoned by NTRs assemble into NPCs ^39,40,57–59^.

We propose that Ran-regulated chaperoning of peripheral Nups by NTRs can foster or impair their association with the NPC core scaffold in a compartment-specific manner. This would ensure that dependent on the RanGTP level, one set of peripheral Nups is dissociated from NTRs and available for association with NPCs, while the other set is sequestered by NTR binding. In our experiments, the nuclear peripheral Nups Nup153 and Mgtor localized to AL in embryos when the levels of cytoplasmic RanGTP were increased (Figure 6A-C).

Elevated levels of RanGTP in the cytoplasm, imitating the nuclear environment, likely cause the disassembly of nuclear import complexes containing these nuclear peripheral nucleoporins, thereby permitting their aberrant association with the NPC’s core scaffold in the cytoplasm. Likewise, cytoplasmic peripheral nucleoporins are chaperoned in export complexes while in the nucleus and are made available for binding to NPCs upon RanGTP hydrolysis in the cytoplasm. Thus, we propose that Ran regulates NPC asymmetry via NTR-mediated enrichment of peripheral Nups in distinct compartments, and by compartment-specific chaperoning (Figure 6D).

Our model does not directly explain the origin of NPCs with an asymmetric core scaffold, in which the number of Y-complex rings differs between NR and CR, as observed in *D. discoideum* ^48^, the alga *C. reinhardtii* ^49^ and a subset of *S. cerevisiae* NPCs ^45^. However, it is conceivable that peripheral Nup binding and stacking interactions between Y-complexes are interdependent ^12,13,23^.

### NPC conversions

An NPC core scaffold that allows a reversible attachment of any peripheral nucleoporin on both sides would facilitate remodelling of NPCs after transitions between compartments. NPC conversion has been suggested for AL insertion ^17,18^, but conceivably could also occur between nucleoplasmic NPCs and NE-NPCs.

Our model postulates that an exchange of peripheral Nups occurs upon exposure to a different subcellular environment (Figure 6E). This would require a sufficiently high turnover frequency of peripheral Nups at the NPC, which is indeed consistent with earlier fluorescence recovery after photobleaching (FRAP) measurements on nuclei from mammalian cells and yeast. In these studies, peripheral Nups were highly mobile, while inner ring and Y-complex components appeared more static ^60,61^.

Using the Ran gradient as a key regulator of NPC asymmetry provides an elegant mechanism to convert symmetric cNPCs to asymmetric NPCs during AL insertion. Our earlier work had suggested that insertion events in *Drosophila* syncytial blastoderm embryos are frequently accompanied by openings of the NE towards the inserting AL, which would expand the nucleoplasmic RanGTP phase ^17^. It is conceivable that exposure to the RanGTP environment during AL insertion allows nuclear export receptors to bind peripheral cytoplasmic Nups on the prospective nuclear ring (Figure 6E). This would trigger their detachment, while peripheral nuclear nucleoporins that are enriched in the nucleoplasm (Figure S1A) could assemble, making NPC remodelling and insertion concomitant events (Figure 6E).

Nucleoplasmic NPCs might result from rearranging the original NE as a consequence of AL insertion ^17^. We indeed observed a potential intermediate of such a conversion in *Drosophila*, where we trapped an individual nNPC in a membrane topology suggesting a preceding insertion event (Figure S4B). In template matching, this nNPC behaved similar to an NE-NPC with an asymmetric peripheral Nup configuration. Although this observation is anecdotal, it is in line with a model where remodelling of the peripheral NPC configuration is subsequent to the relocation into a different compartment. Fully resolving the detailed mechanism of NPC conversions requires future studies, but our present work suggests that no predefined directionality needs to be observed during NPC relocations.

Taken together, our findings emphasize that protein complex composition is not only determined by gene expression, assembly pathways or posttranslational modifications, but also by the cellular environment. This enables dynamic addition or removal of subunits, depending on where in the cell the complex dwells, without the need for degradation, protein synthesis or de novo complex formation. Thus, local chaperone and transport activity may support the establishment and persistence of dedicated complexes. The localization-dependent regulation of complex composition highlights the importance of studying biological structures *in situ*.

## Acknowledgements

We thank Angela Burkart for maintaining *Drosophila* stocks, Birgit Wolf for fly food preparation, and all the members of the Beck laboratory for discussion and advice. We thank Janina Baumbach for construct design to generate transgenic flies and to express recombinant protein to generate the anti-*Dm*Nup358 antibody. We are grateful to our colleagues V. Doye, A. Ephrussi and Y. Shevelyov for fly stocks and reagents. Stocks from the Bloomington Stock Center (NIH P40OD018537) were used in this study. We thank BestGene Inc. (Chino Hills, CA 91709, USA) for *Drosophila* transgenesis. We are grateful to ImmunoGlobe Antikörpertechnik GmbH (97267 Himmelstadt, Germany) for the production of the polyclonal anti-*Dm*Nup358 antibody. We thank Sonja Welsch, Mark Linder, Simone Prinz and Susann Kaltwasser (Central Electron Microscopy Facility, MPI of Biophysics) for assistance with cryo-EM sample preparation and data acquisition. We thank Özkan Yildiz, Juan F. Castillo Hernandez, Thomas Hoffmann, and the Max Planck Computing and Data Facility for computational resources. We acknowledge support from the Imaging Facility of the MPI of Biophysics and MPI of Brain Research. We acknowledge support from the EMBL Electron Microscopy and Protein Purification and Expression Core facilities. We thank Stefanie Böhm, Beata Turoňová, Yannick Schwab and Sonja Welsch for critical reading of the manuscript. J.S. was supported by EnABLE research network. R.T. was supported by an EMBO-long term fellowship (ALTF 170-2019) and the Osamu Hayaishi Memorial Scholarship for Study Abroad from the Japanese Biochemical Society. S.K. was supported by the International Max Planck Research School for Molecular and Cellular Life Sciences. M.Be. acknowledges funding from the Max Planck Society and from the European Research Council (724379-Complex Assembly and 101054823-NPCvalve).

## Author contributions

Conceptualization, J.S., M.Be., B.H.; Methodology, J.S., S.K., R.T., M.A.K., J.M.P., F.W., M.Bö., M.Be., B.H.; Investigation, J.S., M.Bö., S.K., A.B., R.T., M.A.K., V.P., E.K., F.W., B.H.; Formal Analysis, J.S., M.Bö., R.T., B.H.; Visualization, J.S., B.H.; Writing – Original Draft, J.S., M.Be., B.H.; Writing – Review & Editing, J.S., M.Bö., S.K., R.T., M.A.K., F.W., M.Be., B.H.; Project Administration, M.Be.; Supervision, J.M.P, M.Be., B.H.; Funding Acquisition, J.M.P., M.Be.

## Methods

### Fly husbandry and strains

*Drosophila melanogaster* fly stocks were maintained at 22°C on standard cornmeal agar in round-bottom vials. For live imaging or fixation of egg chambers, young (<7 d) females and roughly half as many male flies were transferred to a fresh vial supplemented with yeast paste 24-48 h before the experiment. Embryo collections for the syncytial blastoderm stage were performed on apple-juice plates supplemented with fresh yeast paste for 0-3 h at 22 °C. The following fly lines were used in this study: *w*wg^Sp-^*^1^*/CyO;P{mGFP-Nup107.K}9.1* ^62^(BDSC-35514); *w*wg^Sp-^*^1^*/CyO;P{mRFP-Nup107.K}7.1* ^62^(BDSC-35517); *y1 w*; P{Mgtor.mCherry}2* ^63^ (BDSC-93153); *w*P{w[+mC]=UASp-RFP.KDEL}10/TM3,Sb[1]* (BDSC-30909) (gift from A. Ephrussi); *w[1118];; emeraldGFP::Nup358,PBac{y[+mDint2]=vas-Cas9}VK00027* ^21^*. UASp-RFP.KDEL* in blastoderm embryos was maternally expressed by an insertion of *P{oskar-Gal4}* (FBtp0083699) on the second chromosome ^64^ (gift from A. Ephrussi)*. w[1118]* (BDSC-5905) was used as wild type control.

### Generation of fluorescently tagged transgenic flies

CRISPR-induced homology-directed repair ^65^ was used to endogenously tag nucleoporins with the monomeric GFP variant mEmerald, mCherry or the two part of the split-Venus system, VN (aa 1-173) or VC (aa 155-236) ^28^. Double strand breaks were introduced by injection of a pU6-BbsI-chiRNA vector (Addgene #45946) carrying a complementary guide RNAs (see Table S1 for details) designed via the flyCRISPR target finder tool (http://targetfinder.flycrispr.neuro.brown.edu). A donor repair template (backbone C-terminal tagging: pHD-DsRed - Addgene #51434; backbone N-terminal tagging: pUC19-Addgene #50005) carrying the tag as well as a loxP-DsRed-loxP eye marker cassette flanked by 600-1000 bp homology regions up- and downstream of the start codons (for N-terminally tagged mCherry::Nup358, GFP::Nup98, GFP::Nup133 and mCherry::Nup153 flies) or the stop codons (for C-terminally tagged Nup358::VN, Nup358::VC, Nup358::GFP and Nup214::GFP flies) was co-injected to produce endogenously tagged fusion proteins. In this repair template, the CRISPR targeting sequence was mutated. Depending on the modified chromosome either *y[1] sc[*] v[1] sev[21]; P{y[+t7.7] v[+t1.8]=nanos-Cas9.R}attP40* (BDSC-78781) ^66^, *w[1118]; PBac{y[+mDint2] GFP[E.3xP3]=vas-Cas9}VK00027* flies (BDSC-51324) ^65^, or *yw; nos-Cas9(II-attp-40)* ^67^ *flies* were injected. See Table S1 for line-specific information.

Transgenic individuals were identified by the presence of the eye marker cassette. After removal of the eye marker cassette by Cre-mediated recombination, homozygous male and female transgenic flies were viable, fertile and showed no obvious phenotypes for all lines except for Nup358::VN transgenic flies. Homozygous Nup358::VN females were viable but sterile and only a heterozygous stock could be maintained. All insertions were characterized by genomic PCR and sequencing.

### Expression of recombinant Ran proteins

The gene encoding full-length *Hs*Ran (UniProt ID: P62826) was cloned into the pGEX-6P-1 expression vector. The GST tag was subsequently replaced with a 6xHis-tag for histidine affinity purification. The Q69L point mutation was introduced to the construct by site-directed mutagenesis.

Ran proteins were expressed in *E. coli* BL21-CodonPlus(DE3)-RIL strain (Agilent). Transformed cells were grown at 37 °C in LB medium containing 100 µg/ml Ampicillin up to an OD_600_ of 0.6, and then 0.5 mM isopropyl-β-D-thiogalactopyranoside was added to induce the protein expression. Cells were subsequently cultured at 18 °C overnight and collected by centrifugation (4,700 rcf, 20 min, RT). Cell pellets were stored at −80 °C. For purification, the frozen pellet was resuspended in a buffer containing 20 mM Tris-HCl (pH 8.0), 200 mM NaCl, 5 mM imidazole (pH 8.0), 3 mM β-mercaptoethanol, 4 mM MgCl_2_, 1 mM benzamidine, 1 mM phenylmethylsulfonyl fluoride, complete EDTA-free protein inhibitor cocktail (Roche) and DNAse I (Roche). The cell suspension was filtered using a nylon sieve (Roth, TA91.1), and was applied to a microfluidizer for cell lysis. After clearing the crude lysate by ultracentrifugation (185,500 rcf, 1 h, 4 °C), the supernatant was mixed with Ni-NTA agarose resin (Macherey-Nagel) and incubated for 1.5 h at 4 °C. The resin was washed with a buffer containing 10 mM Tris-HCl (pH 8.0), 400 mM NaCl, 10 mM imidazole (pH 8.0), and 0.2 mM TCEP, and Ran proteins were subsequently eluted from the resin with a buffer containing 10 mM Tris-HCl (pH 8.0), 100 mM NaCl, 200 mM imidazole (pH 8.0), and 0.2 mM TCEP. The eluted sample was mixed with HRV-3C protease (Merck, SAE0045) to cleave off the 6xHis-tag, and was dialyzed in a buffer containing 10 mM Tris-HCl (pH 8.0) and 100 mM NaCl overnight to remove the imidazole. The dialyzed sample was again loaded onto Ni-NTA agarose resin (Macherey-Nagel) to remove the 6xHis-tag, HRV-3C protease, and non-specifically bound contaminants. The flow-through fraction was then concentrated using an Amicon Ultra filter (Merck, molecular weight cut-off 10 kDa), and was further purified by size-exclusion chromatography on a HiLoad 16/600 Superdex 200 pg column (Cytiva), in a buffer containing 15 mM Tris-HCl (pH 8.0), 150 mM NaCl, 1 mM MgCl_2_, and 0.2 mM TCEP. The peak fractions were concentrated to approximately 5 mg/ml using an Amicon Ultra filter (Merck, molecular weight cut-off 10 kDa). The sample was subsequently supplemented with glycerol at a concentration of 10 %_v/v_, aliquoted, and stored at −80 °C until use.

### Generation of anti-*Dm*Nup358 antibody

Anti-*Dm*Nup358 polyclonal antibodies were generated in rabbits by immunization against a recombinant portion of the *Dm*Nup358 protein (aa 1770-1918). To generate a recombinant portion of the protein, the *Dm*Nup358 sequence was amplified by PCR from cDNA (pOT-LD43045, BDGP) and cloned into pGEX_6P1 to express a recombinant protein with an N-terminal GST and a C-terminal GFP tag. The plasmid was transformed into *E. coli* BL21-CodonPlus (DE3)-RIL strain (Agilent) and transformed cells were grown in an overnight pre-culture in LB medium containing 50 µg/ml chloramphenicol and 100 µg/ml carbenicillin.

Large scale expression was induced in TB-FB medium, 1.5 % lactose, 0.05 % glucose, 2 mM MgSO_4_, 50 µg/ml chloramphenicol, 100 µg/ml carbenicillin and 100 µM ZnCl_2_. Cells were grown to an OD_600_ of 0.6-0.8 at 37 °C and subsequently cultured at 18 °C overnight and collected by centrifugation (4,700 rcf, 20 min, RT). Cell pellets were stored at −80 °C. For lysis, the frozen pellet was re-suspended in a buffer containing 50 mM Tris-HCl (pH 7.4), 500 mM NaCl, 1 mM DTT, 1 mM Benzamidin, 100 µM ZnCl2, 1 mM MgCl_2_, 1 mM phenylmethylsulfonyl fluoride, complete EDTA-free protein inhibitor cocktail (Roche) and DNAseI (Roche). The cell suspension was filtered using a nylon sieve (Roth, TA91.1), and was applied to a microfluidizer. After clearing the crude lysate by ultracentrifugation (185,500 rcf, 1 h, 4 °C), the supernatants were filtered (0.45 µm) and loaded onto a GSTrap4B (5ml, Cytiva) for affinity purification. After washing with the same buffer, proteins were eluted in 50mM Tris-HCl (pH 7.4), 500 mM NaCl, 1 mM DTT, 30 mM Glutathion. Eluted protein was concentrated (Amicon Ultra filter, Merck, molecular weight cut-off 50 kDa) and desalted with a PD10 column (GE #17085101) before anion-exchange. The desalted protein was subsequently loaded in buffer A (50 mM Tris-MES, pH 7.4, 0.2 mM TCEP) onto a Resource Q (1ml, Cytvia) for anion exchange chromatography, washed with 5CV of buffer A and eluted with buffer A containing 2M NaCl with a linear gradient up to 1M NaCl for 40CV. Eluted pooled fractions were concentrated using an Amicon Ultra filter (Merck, molecular weight cut-off 50 kDa) and applied for size exclusion chromatography on a HiLoad 16/600 Superdex 200pg (Cytvia) in PBS, 100 mM NaCl, pH 7.4, 0.2 mM TCEP. Eluted proteins were frozen in liquid nitrogen and stored at −80 °C.

To generate antisera, a rabbit was immunized with 242 µg of recombinant protein antigen 1:1 with Montanide ISA206 (ImmunoGlobe GmbH). After bleeding the serum was pre-cleared on an NHS-activated sepharose affinity matrix containing GST-GFP (expressed and purified similar to the recombinant portion of Nup358 used for immunization) and subsequently subjected to fractional purification (first against GST-GFP, followed by affinity purification with the immobilized antigen). Antibodies were frozen in Tris-HCl, 0.002 % NaN_3_, sterile filtered and stored with 25% glycerol at −80 °C.

### Live imaging of *Drosophila* embryos and egg chambers

Ovaries from females of the desired genotype were dissected in BRB80 buffer (80 mM PIPES pH 6.9, 1 mM EGTA, 2 mM MgCl_2_) and mounted onto coverslips in a drop of Schneider’s S2 medium supplemented with 10 % fetal calf serum and insulin (200 µg/ml). Adjacent to the ovaries containing buffer, a drop of Voltalef 10S oil (VWR) was placed onto the coverslip and individual ovarioles were pulled into the oil with fine tungsten needles ^68^. For embryo live-imaging staged syncytial blastoderm embryos were dechorionated, aligned and glued onto a coverslip and overlaid with Halocarbon oil before imaging ^69^. Embryos and egg chambers were imaged at RT on an inverted Stellaris 5 confocal microscope (Leica) with a 63x/1.4 NA oil immersion objective.

### *Drosophila* embryo injections

Staged syncytial blastoderm embryos of the respective genotype were dechorionated, aligned and glued onto a coverslip. Embryos were dried on silica for 8 min, overlaid with Halocarbon oil and injected together with WGA-Alexa647 (100 μg/ml). GTPγS (10 mM) was injected in water. *Hs*Ran and *Hs*RanQ69L (both at 6 mg/ml) were injected in a buffer containing 15 mM Tris-HCl (pH 8.0), 150 mM NaCl, 1 mM MgCl_2_, and 0.2 mM TCEP. Imaging was performed immediately (<5 min) subsequent to injection.

### Immunofluorescence of *Drosophila* embryos and egg chambers

Staged syncytial blastoderm embryos of the respective genotype were dechorionated and fixed in equal volumes of heptane and 4 % Formaldehyde in PBS for 20 min rotating. Embryos were devitellinized by Methanol, washed with PBS, 0.1 % Triton X-100 and blocked for 1 h in blocking buffer (PBS, 0.1 % Triton X-100, 10 % Normal Goat Serum) at RT. Primary antibodies were applied in blocking buffer overnight at 4 °C. After washing in PBS, 0.1 % Triton X-100, secondary antibodies (Alexa conjugates, Life Technologies) were applied in blocking buffer (1:500) for 2 h at RT. Samples were mounted after 3 times 10 min washes (PBS, 0.1 % Triton X-100) onto coverslips using Vectashield and kept at 4 °C until imaging acquisition.

For immunostainings of egg chambers, ovaries from ∼5 well fed females of the respective genotype were dissected in PBS, 0.05 % Triton X-100 and immediately fixed in 4 % Formaldehyde in PBS for 30 min at RT. After permeabilization in PBS, 1 % TritonX-100 for 1 h at RT, ovaries were incubated in blocking solution (PBS, 0.1 %Triton X-100, 10 % Normal Goat Serum) for 1 h at RT. Incubation with primary and secondary antibodies was similar to embryo immunostainings. Upon mounting in Vectashield, ovaries were further separated into ovarioles or single egg chambers and samples were kept at 4 °C until imaging. Image acquisition of fixed embryo and egg chamber samples was performed on an inverted Stellaris 5 confocal microscope (Leica) with a 63x/1.4 NA oil immersion objective. The following antibodies were used in this study: rabbit anti-*Dm*ELYS (1:1000) ^70^; rabbit anti-*Dm*Nup358 (this study).

### Quantification of relative nucleoporin abundance

Regions of interests (ROIs) were hand selected in Fiji ^71^ for NE, AL and cytoplasmic regions devoid of Nup fluorescence in the composite images recorded from living syncytial blastoderm embryos expressing pairs of fluorescently tagged Nups. Subsequently the mean fluorescence intensity was measured for each ROI on the separated channel. The mean intensities for AL and NE were divided by the mean of the respective cytoplasmic intensity in each image to normalize for background fluorescence. The ratio of these normalized intensities was calculated for each pair of Nups for NE and AL ROIs. To compare between different images and embryos, all ratios were normalized to the mean ratio at the NE for each Nup pair.

### In-section correlative light and electron microscopy (CLEM)

CLEM analysis was done as previously described ^21^. Shortly, dissected ovaries or staged embryos of the respective genotype were high-pressure frozen (HPM010, AbraFluid) in Schneider’s S2 medium containing 20 % Ficoll (70 kDa) as a cryoprotectant. The samples were freeze-substituted (EM-AFS2, Leica Microsystems) with 0.1 % Uranyl Acetate (UA) in acetone at −90 °C for 72 h. The temperature was then raised to −45°C at 3.5 °C/h and samples were further incubated for 5 h. After rinsing in acetone, the samples were infiltrated in Lowicryl HM20 resin, while raising the temperature to −25 °C and left to polymerize under UV light for 48 h at −25 °C and for further 9 h while the temperature was gradually raised to 20 °C (5 °C/h). Thick sections (300 nm) were cut from the polymerized resin block and picked up on carbon coated mesh grids (Plano, S160). The fluorescence microscopy imaging of the sections was carried out as previously described ^72^ using a widefield fluorescence microscope (Olympus IX81 with MT20 illumination system) equipped with an Olympus PlanApo 100X 1.40 NA oil immersion objective.

10 nm protein A gold (UMC Utrecht, Cell Microscopy Core) was added to the sections as (tomographic) fiducial markers, before post-staining with uranyl acetate and lead citrate. (Dual-axis) tilt series of the areas of interest were acquired over a −60° to +60° tilt range (1° increment) using a FEI Tecnai F30 TEM (300kV, Gatan OneView camera; final pixel size 1.55 nm) and tomograms were reconstructed using the software package IMOD ^73^.

Correlation between light and electron micrographs was carried out with the plugin ec-CLEM ^74^ of the software platform Icy ^75^. The coordinates of pairs of corresponding features in the two imaging modalities were used to calculate a similarity transform (translation/rotation/scaling), which allowed to map the coordinates of the fluorescent spots of interest and to overlay them on the electron tomogram.

### *Drosophila* ovary cell preparation for cryo-ET

Small cells from *Drosophila* ovaries were isolated by dissection, tissue dissociation and filtration as previously described ^29^. Shortly, about 15 ovaries were prepared from female flies into Schneider’s S2 medium supplemented with 10 % fetal bovine serum, 50 U/ml Penicillin, 50 µg/ml Streptomycin and 0.25 mg/ml fungizone in a 2 ml Eppendorf tube.

Ovaries were washed three times in 700 µl calcium-free phosphate-buffered saline and sedimented by gravity in between. For tissue dissociation, ovaries were incubated in 0.5 % Trypsin in PBS for 15 min at room temperature, shaking at 900 rpm. Ovaries were sedimented by gravity, and the supernatant containing dissociated cells was filtered through a 40 µm cell strainer into a Falcon tube. 700 µl S2 medium was used to wash the remaining cells through the cell strainer. The filtrate was transferred to a fresh 1.5 ml Eppendorf tube and cells were sedimented for 7 min at 5000 rcf. After removal of the supernatant, the pelleted cells were washed with 1.4 ml sterile S2 medium and sedimented again for 7 min at 5000 rcf. Trypsinization, filtration of the supernatant and washing of the filtrate was repeated twice with the remaining tissue. The final cell pellet containing cells from all three dissociation steps was resuspended in 20-40 µl fresh S2 medium and directly subjected to plunge freezing.

3 µl cell suspension was applied on glow-discharged (Pelco easiGlow) R1/4 SiO_2_ gold 200 mesh grids (Quantifoil), blotted for 10 sec from the backside and plunge-frozen on a Leica EM GP2 plunger at 23 °C, 70 % relative humidity. Up to 5 %_v/v_ glycerol was added to the cell suspension directly before the freezing session to improve vitrification. Due to unreliable blotting, 3 µl of medium of the appropriate glycerol concentration was applied to the backside of the grid before freezing. Data used in this study came from four separate sample preparation sessions.

Grids were assembled into AutoGrid cartridges and loaded into an Aquilos 1 cryo-FIB-SEM (Thermo Scientific) to prepare lamellae similar to established protocols ^76,77^. In brief, the sample was coated with an organometallic layer with a Gas Injection System for 10-15 sec.

Subsequently, an inorganic platinum layer was applied by sputter coating at 1 kV, 10 mA, 10 Pa for 20-25 sec. A gallium beam at 30 kV was used to prepare lamellae in a stepwise fashion, reducing the beam current from 1 nA (5 µm thickness), to 500 pA (3 µm thickness), to 300 pA (1.2 µm thickness) to 100 pA (700 nm thickness). Rough milling steps were carried out by SerialFIB ^77^. Final polishing to the target thickness of 150-200 nm was done with 50 pA current. Lamella milling was monitored by SEM at 25 pA at 2-10 kV. A final inorganic platinum layer was applied by sputter coating at 1 kV, 10 mA, 10 Pa for 3 sec before unloading.

### Yeast strains and growth conditions

The *MATa, ura3Δ0, leu2Δ0, his3Δ1, met15Δ0, Δmot2::kanMX4, NUP188::EGFP::HIS3MX6, NUP170::mars::hphNT1*, called Not4Δ Nup188::GFP Nup170::mars in this manuscript, were stored at cryo-stocks in YPD with 50 % glycerol and cultured fresh for every sample preparation. Shortly, YPD plates were streaked from the cryostocks and colonies were grown at 30 °C for 3-6 days. Single colonies were used to inoculate 5 ml of synthetic minimal medium supplemented with 2 % glucose, grown at 30 °C at 220 rpm in shaker overnight before cryo-ET sample preparation.

### Yeast sample preparation for cryo-ET and cryo-correlative imaging

Liquid yeast cultures were grown to OD_600_ of 0.3-0.8 for vitrification. 3.5 µl cell suspension applied on glow-discharged (Pelco easiGlow) R1/4 SiO_2_ gold 200 mesh grids (Quantifoil), blotted for 2 sec from the backside and was plunge-frozen on a Leica EM GP2 plunger at 20 °C, 90 % relative humidity. Occasionally, 1 µl media was added to that back of the grid to help with blotting. Data used in this study came from four separate sample preparation sessions.

Lamellae were prepared on an Aquilos 1 or Aquilos 2 cryo-FIB-SEM (Thermo Scientific) as described for the *Drosophila* samples, but an additional correlation step was used to improve targeting of NPC clusters (Figure 5A): AutoGrid cartridges were inserted into the shuttle of the cryo light microscope (Stellaris SP8 cryoCLEM, Leica), which was further fastened into the Aquilos Leica CLEM Shuttle for lamella preparation. In the first cryo-FIB-SEM session, lamellae were milled to roughly 1 µm thickness. Subsequently, the light microscope shuttle including the two rough milled grids was transferred to the cryo light microscope to record confocal stacks of the lamellae in the red (excitation: 552 nm, detection: 596-703 nm) and green (excitation: 488 nm, detection: 495-522 nm) channel, as well as in transmitted mode (T-PMT), through the HC PL Apo 50X dry objective (NA 0.9).

Maximum intensity projections were created in ImageJ/Fiji ^71^ and used to select lamellae to be polished, and to guide TEM acquisition. Correlation between the SEM and the FM imaging was performed manually based on the grid overviews and the consistent orientation ensured by keeping the grids inside the cryo light microscope shuttle through the procedure. Lamella with NPC clusters were polished in a second cryo-FIB-SEM session, in which contamination with ice resulting from the transfer steps could be removed without problems due to the pre-existing protective platinum layer.

### Cryo-ET data acquisition and tomogram reconstruction

Low-dose tilt series were acquired on a Titan Krios TEM, operated at 300 kV in EFTEM mode, using a dose-symmetric tilt scheme with a target dose of 150 e^-^/A^2^ around the pretilt of the lamella in 2° increments grouped by 2. The *Drosophila* data set was recorded with a target defocus range of −1 to −5 µm at a pixel size of 2.414 Å. The *S. cerevisiae* data set was recorded with a target defocus range of −2 to −5 µm at a pixel size of 2.176 Å. Each projection was recorded as 10 fractions that were motion-corrected in SerialEM ^78^ on-the-fly.

A total of 466 tilt series (TS) were acquired in the *Drosophila* sample. Excluding TS that were of insufficient quality or did not contain NPCs resulted in 111 TS that were used further. 467 TS were acquired in the yeast sample. Excluding TS that were of insufficient quality or did not contain nNPCs resulted in 40 TS that were used further.

Projection images were filtered by cumulative electron dose ^79^ in Matlab as previously described ^80^ and gctf v1.06 ^81^ was used for defocus estimation. Tilt images of poor quality or unsuccessful defocus estimation were assessed visually and low-quality tilts were discarded. The resulting cleaned dose-filtered TS were aligned by patch tracking using either IMOD ^73^ or AreTomo ^82^ and reconstructed as bin 4 tomograms. IMOD’s SIRT-like filtering and deconvolution implemented in MemBrain ^83^ were applied for visualization (Figure 3A, Figure 4A, Figure S4Ai and Bi, Fig 5B). NovaCTF ^84^ was used to generate 3D CTF-corrected tomograms. Further details are to be found in Table S2 (*Drosophila*) and Table S3 (yeast).

### Membrane segmentation

Membranes were automatically segmented using MemBrain-Seg with the model version MemBrain_seg_v10_alpha.ckpt ^83^. Resulting segmentations were manually cleaned from wrongly assigned membranes in UCSF ChimeraX ^85^. Segmentations were visualized in ChimeraX (Figure 3A, Figure 4C-D, Figure S4Aii-iii and Bii-iii, Figure 5F).

### Subtomogram averaging

NPCs were manually picked during visual inspection of the tomograms. Only tomograms without apparent defects were used for subtomogram averaging (STA).

Initial orientation was assigned by the compartment in the case of NE-NPCs, with the top of the average assigned towards the cytoplasm. For NPCs outside the NE, the orientation was assigned based on the closeness of the compartment: For nNPCs, the face closer to the cytoplasm was assigned as the top (cytoplasmic face in NE-NPCs), for cNPCs the face closer to the nucleus was assigned as bottom (nuclear face in NE-NPCs). Subtomograms were extracted, aligned and averaged using novaSTA ^86^ as described previously^31,48^. NE-NPCs, cNPCs and nNPCs were processed separately.

Initially, whole NPCs were aligned and averaged with imposed eightfold symmetry at bin 8. Based on this initial average, the positions of asymmetric units were extracted and aligned at bin 4. After aligning the asymmetric units, alignment was focused on the CR, IR and NR, as well as NB in case of the *Drosophila* NE-NPC. Particles outside the lamella or misaligned with respect to the membrane were manually discarded using ArtiaX ^87^ in ChimeraX. Particles were mapped back into the tomographic volume for visualization using ArtiaX in ChimeraX (Figure 3A, Figure 5B).

Masks for averaging and for FSC calculation were generated using ChimeraX, Dynamo ^88^, cryoCAT [Turoňová, B. https://github.com/turonova/cryoCAT] or Relion 3.1.3 ^89^. FSC curves represent phase-randomization corrected values calculated by Relion 3.1.3 (Figure S3A, Figure S5A).

To generate the composite map of the whole NPC, the final averages of the individual rings were fitted into the map of the asymmetric unit and assembled into an eightfold symmetric architecture based on the coordinates used for asymmetric unit extraction (Figure 3B-D, Figure S3B, Figure S5B,Figure 5C-D). Further details are to be found in Table S2 (*Drosophila*) and Table S3 (yeast).

### NPC diameter measurements

To estimate diameters of the NPC rings, the coordinates from STA were used. STA maps were aligned to a common orientation using ChimeraX and cryoCAT and this alignment was transferred to individual particles. The diameters were then measured as pairwise distances between opposite subunits at specified points (Figure S3B and Figure S5B) from the coordinates using cryoCAT. Finally, the average of all pairwise distances per NPC is calculated (Figure S3C, Figure S5C).

### Three-dimensional template matching

Three-dimensional template matching was performed with GPU Accelerated Python Stopgap for Template Matching (GAPSTOP^TM^) ^34,90^ on bin 4 tomogram. The NE-NPC’s CR and NR subtomogram averages which contained an asymmetric unit including the connections to the neighbouring subunits were used as search templates. The NR and CR templates of each species were brought into the same orientation, low-pass filtered (yeast: 42 Å, *Drosophila*: 32 Å) and a constrained cross-correlation (CCC) score was calculated at an angular sampling of 6 degrees. For visualization, scores volumes were displayed in ChimeraX (Figure 4B, Supplementary Video S1).

To determine the threshold for peak extraction, the mean and standard deviation of the central 300×300×100 of the scores volume, which typically contain no non-biological material, was calculated. Peaks with CCC values that are more than 10 (*Drosophila*) or 8 (yeast) standard deviations higher than the mean and at a distance of 25 voxels from each other were extracted. Pixels located closer than 60 voxels to the edge of the reconstructed volume were excluded to avoid edge artifacts.

The results were manually cleaned by backmapping the detected CR and NR particles on the tomographic volume using ArtiaX in ChimeraX. Only particles inside the lamella volume and consistent with membrane association were retained. In case of overlapping particles within one particle class, the particle with the lower CCC score was removed. Matches were visualized in ChimeraX (Figure 4C-D, Figure S4Aii-iii and Bii-iii, Figure 5F).

### Quantification of template matches per ring

Matches are manually classified by their subcellular location into NR of NE-NPCs (on the inner nuclear membrane), CR of NE-NPCs (on the outer nuclear membrane), outer rings of cNPCs (on other membranes in the cytoplasm), and outer rings of nNPCs (on other membranes in the nucleoplasm).

The particle lists of the NR and CR matches were assigned to outer rings. This was done by tracing neighbouring asymmetric units using the tracing function of cryoCAT as previously used ^91,92^, and grouping traced objects based on their distance and orientation. The number of matches obtained for each outer ring for both NR and CR was counted. For a canonical NPC, a number of 8 asymmetric units would be expected. However, incomplete detection and NPCs that are only partially inside the lamella volume cause lower number of matches detected per outer ring in many cases.

The distribution of the combinations of matches observed in each category was depicted in a bubble chart, where the circle area scales with the number of distinct outer rings exhibiting the given number of matches (Figure 4E, Figure S4C, Figure 5E). An orthogonal distance regression in SciPy ^93^ was used to fit the proportionality of observed #NR/#CR matches.

