## Supplementary Information for "The small GTPase Ran defines Nuclear Pore Complex Asymmetry"

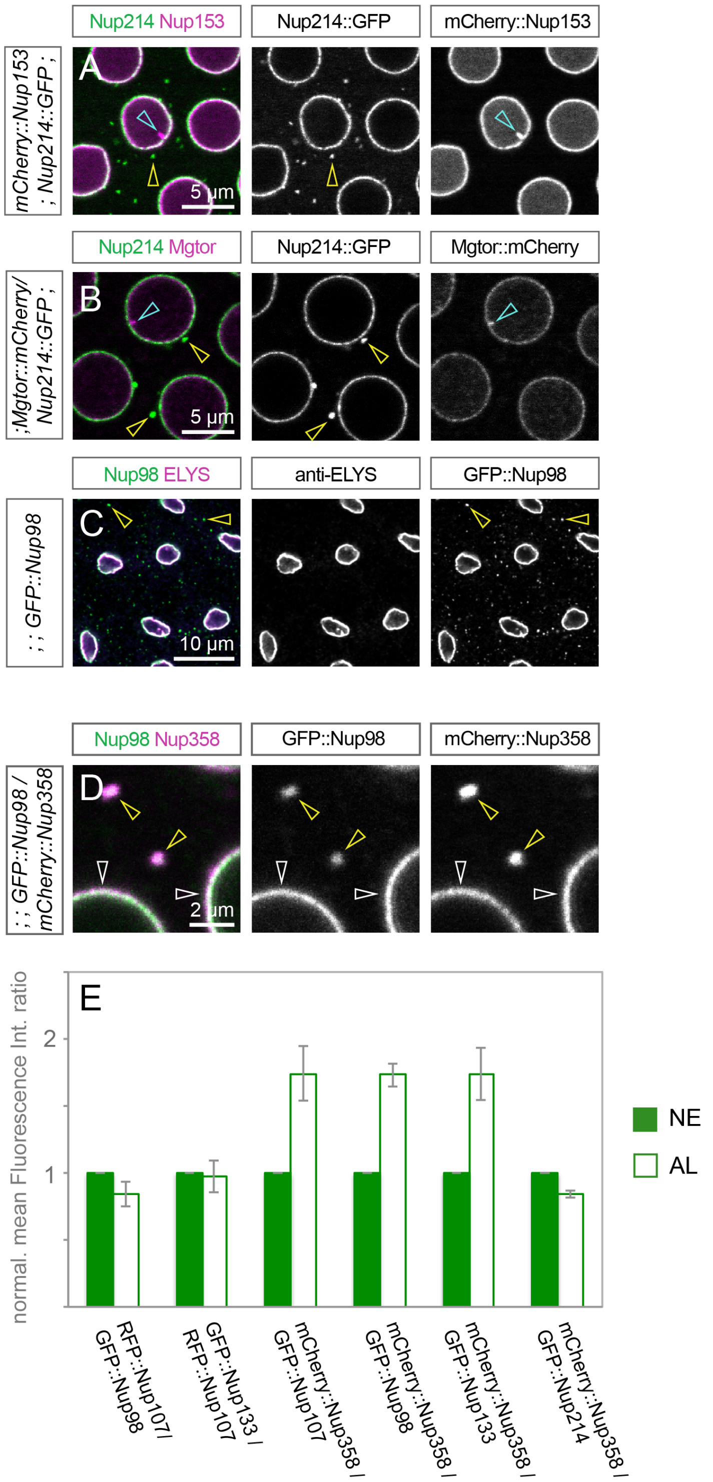


**Figure S1: NPC composition depends on the subcellular context, related to Figure 1**

**A-C** Distinct localization of different nucleoporins in ectopic cytoplasmic and nucleoplasmic NPCs. Confocal images of living (A, B) or fixed (C) *Drosophila* syncytial blastoderm embryos of the indicated genotype. **A, B** The peripheral cytoplasmic Nup214::GFP localizes to cytoplasmic NPCs (yellow arrowheads) but not to nucleoplasmic NPCs (cyan arrowheads). The peripheral nucleoplasmic mCherry::Nup153 (A) and Mgtor::mCherry (B) localize to nucleoplasmic, but not to cytoplasmic NPCs. All Nups are present at the NE. **C** ELYS is absent from NPCs in the cytoplasm, labelled by the NPC core scaffold component GFP::Nup98 (yellow arrowheads). Both proteins are present at the NE. **D** Peripheral cytoplasmic Nups are enriched at cytoplasmic NPCs. **E, D** Peripheral cytoplasmic Nucleoporins are enriched at AL compared to the NE. **D** Confocal image of a *Drosophila* syncytial blastoderm embryo expressing the scaffold component GFP::Nup98 and the peripheral cytoplasmic mCherry::Nup358. Qualitatively, Nup358 appears enriched at AL compared to Nup98 (yellow arrowheads), relative to their distribution at the NE (white arrowheads). **E** Quantification of relative nucleoporin abundances at NPCs at AL and the NE. Mean fluorescence intensities were measured from ROIs at the NE, AL and the cytoplasm for background correction. Ratios of the respective nucleoporin pairs at AL and the NE are plotted, normalized to the NE ratio to compare between genotypes and embryos. Data are presented as means +/- STDV, errors indicate variability over several embryos. The core scaffold components GFP::Nup107, GFP::Nup133 and GFP::Nup98 are equally distributed at the NE and at AL, while mCherry::Nup358 is present at higher levels at AL compared to core scaffold Nups. At AL the ratio of the peripheral cytoplasmic Nups mCherry::Nup358 and GFP::Nup214 is similar.

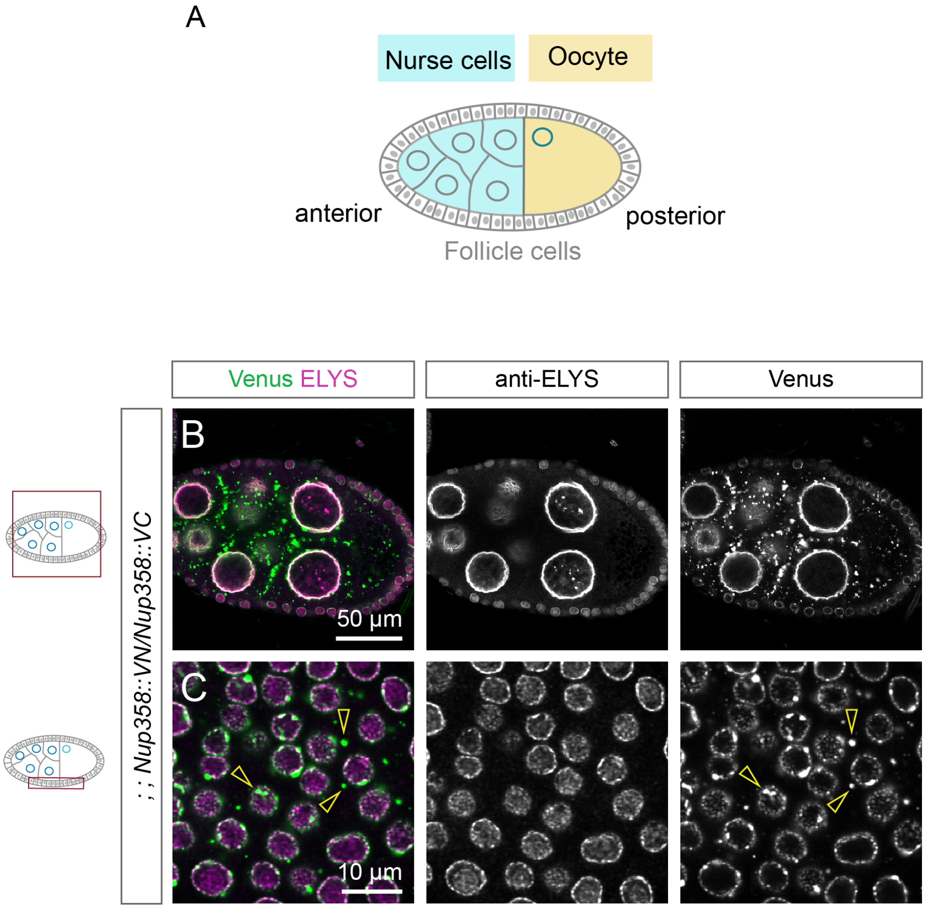


**Figure S2: NPC composition in induced Annulate Lamellae, related to Figure 2**

**A** Schematic representation of a *Drosophila* egg chamber. Germline cells comprise nurse cells and the posteriorly located oocyte. A somatic epithelium of follicle cells surrounds the germline cells. **B-C** Confocal images of a fixed and immunostained *Drosophila* egg chamber dissected from Nup358::VN/Nup358::VC ovaries. The peripheral nucleoplasmic nucleoporin ELYS localizes to the NE, but not to Nup358::VN/Nup358::VC induced AL both in germline cells (B) and in follicle cells (yellow arrowheads) (C).

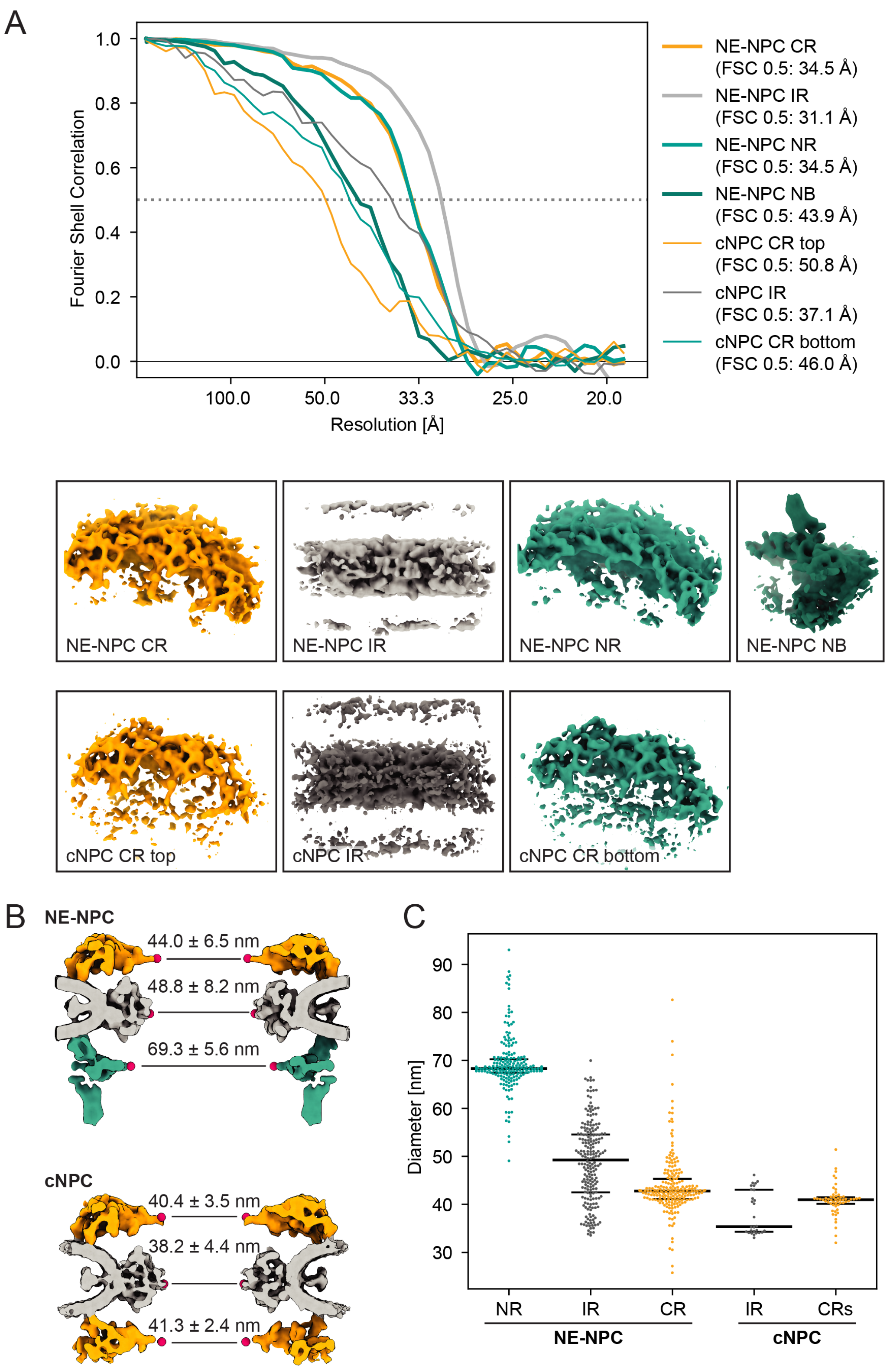


**Figure S3: Resolution of subtomogram averages and diameters of *Drosophila* NPCs, related to Figure 3**

**A** Resolution of subtomogram averages contributing to the composite maps shown in Figure 3. FSC curves of the individual focused subtomogram averages shown in the subpanels below. The resolution cut-off at 0.5 is indicated. Subpanels show the focused averages of the inner ring (IR), the two outer rings (CR and NR) and the nuclear basket (NB) in case of the NE-NPC. Directional picking of the cNPCs results in two separate structures for the cNPC's CRs: CR top and CR bottom are named according to where they are shown in side views of the cNPC. Detailed information on subtomogram averages in Table S2. **B** cNPCs are more constricted than NE-NPCs. Side-cut view of the composite maps with overlay of measurement positions. Diameters of the three rings were measured between the most inward-facing densities of the NE-NPC. Measurement positions in the cNPC were selected consistent with those in the NE-NPC in spite of slight deviations in the map densities. Average diameters (means ± SD) are shown for each ring. **C** Plot of the diameter measurement. Dots indicate measurements on individual rings; the median and upper and lower quartile are indicated by black lines. **A-C:** Sample: Cells isolated from Nup358::VN/Nup358::VC *Drosophila* ovaries.

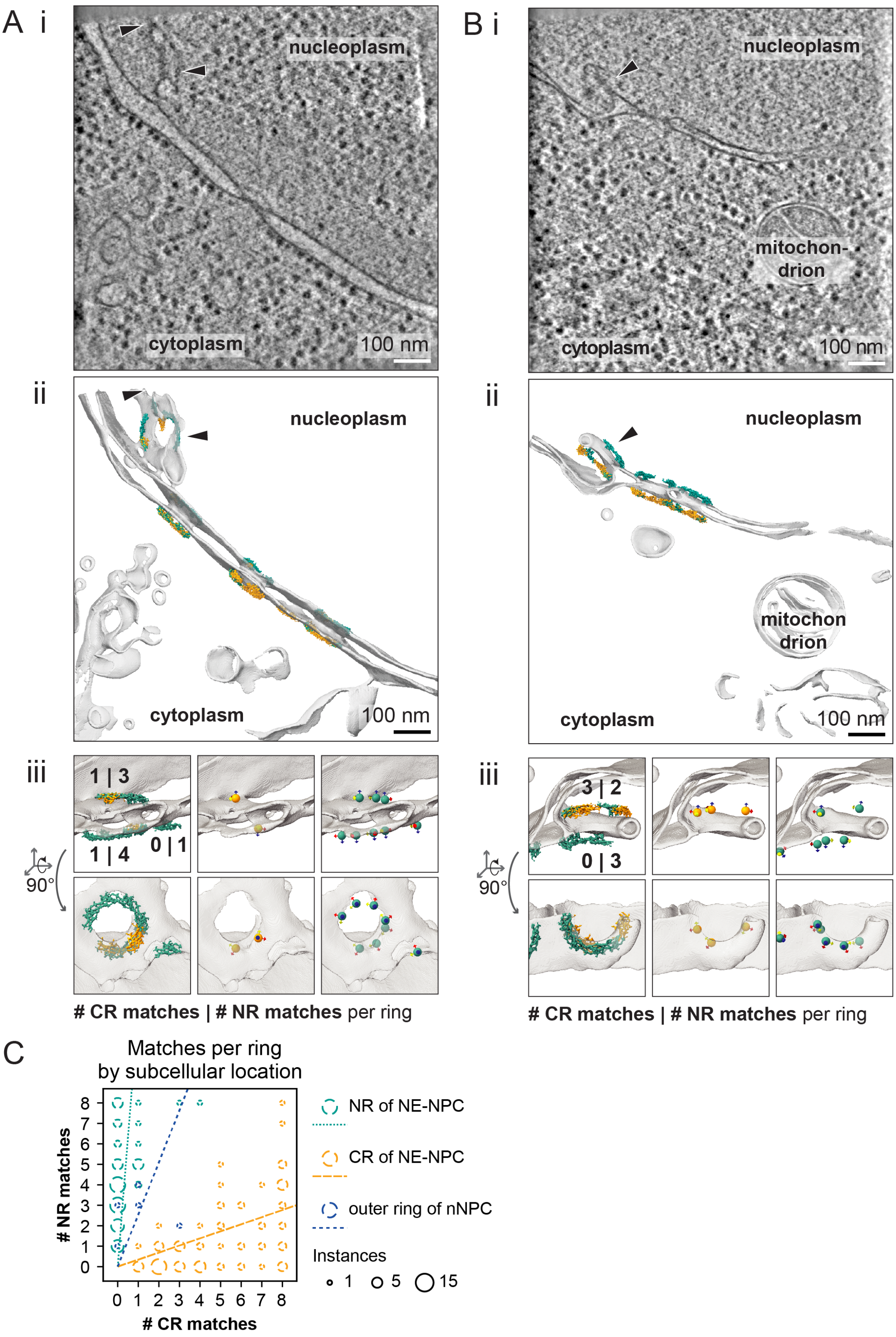


**Figure S4: Template matching suggests that nNPCs in *Drosophila* contain two nuclear rings, related to Figure 4.**

Two tomograms of cells isolated from Nup358::VN/Nup358::VC *Drosophila* ovaries show nucleoplasmic NPCs (arrowheads) that were analysed by using an asymmetric units of the

NE-NPC's CR (orange) and NR (teal) (Figure 3B) as search templates. **A, B** nNPC outer rings are predominantly recognized by the NR template. Slice of a tomogram (subpanel i) and membrane segmentation with mapped back template matches (subpanels ii-iii). Subpanels (iii) show insets of the subvolume containing nNPCs in top and side views. Matches are either indicated by mapped back templates or spherical markers. The number of matches with the two templates is indicated per outer ring, written as #CR matches|#NR matches. **B** The nNPC is embedded in a membrane extending into the nucleoplasm at a position where the NE membrane is highly curved, indicating a preceding remodelling event. The NE-proximal ring of this nNPC exhibits more matches with the CR than the NR template, while the NE-distal outer ring has only matches with the NR template, suggesting yet incomplete conversion of an NE-NPC into an nNPC. **C** nNPC outer rings mostly behave NR-like in template matching. Bubble chart of the matches obtained with the NR template and the CR template per outer ring, grouped by subcellular location. Dashed lines indicate the proportionality of NR vs. CR template matches for each subcellular location. **A-C:** Sample: Cells isolated from Nup358::VN/Nup358::VC *Drosophila* ovaries.

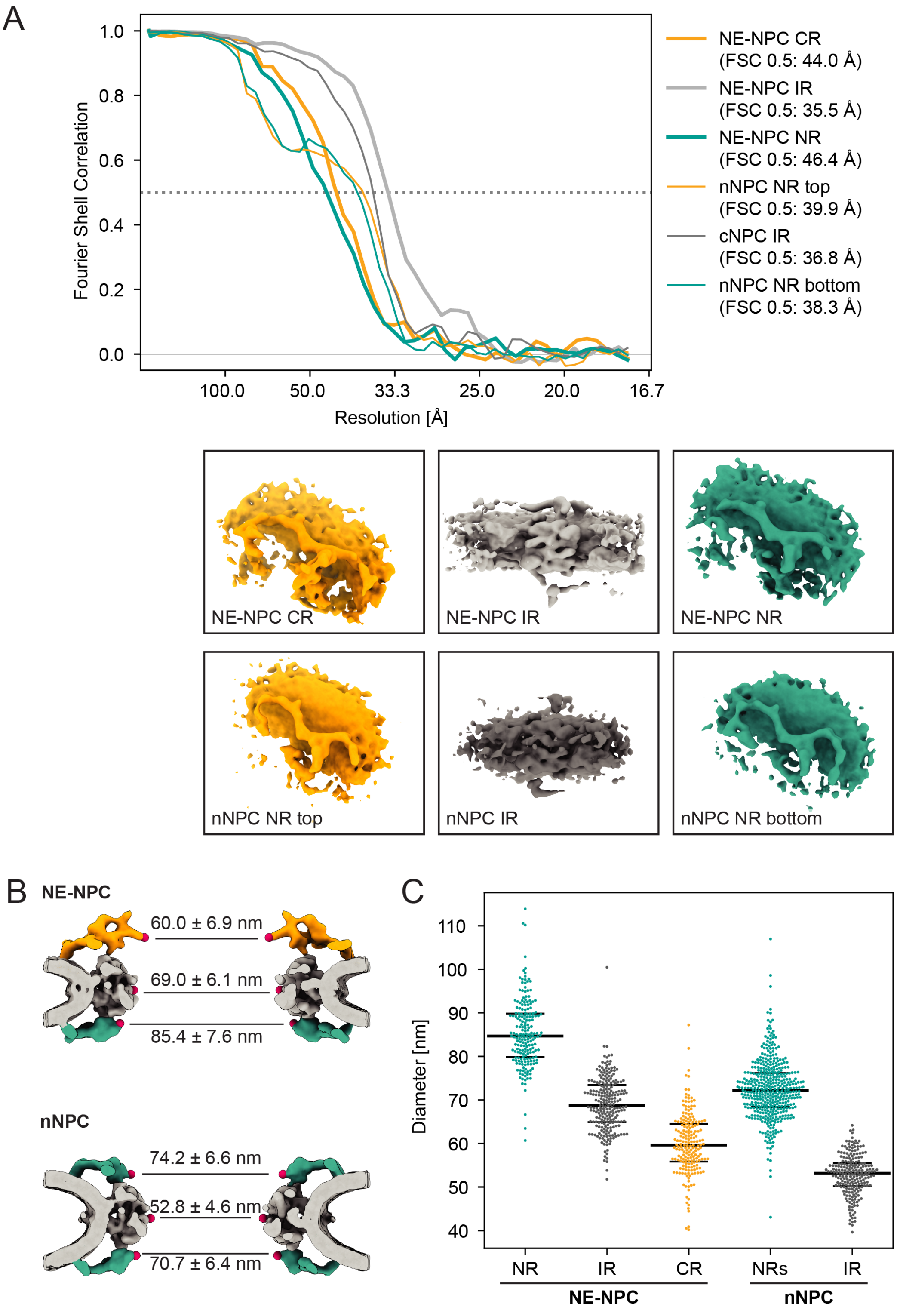


**Figure S5: Resolution of subtomogram averages and diameters of yeast NPCs, related to Figure 5**

**A** Resolution of subtomogram averages contributing to the composite maps shown in Figure 5. FSC curves of the individual focused subtomogram averages. The resolution cut-off at 0.5 is indicated. Subpanels show the focused averages of the inner ring (IR) and the two outer rings (CR and NR). Directional picking of the nNPCs results in two separate structures for the nNPC's NRs: NR top and NR bottom are named according to where they are shown in side views of the nNPC. Detailed information on subtomogram averages in Table S3. **B** nNPCs are more constricted than NE-NPCs. Side-cut view of the composite maps with overlay of measurement positions. Diameters of the three rings were measured between the most inward-facing densities of the NE-NPC. Measurement positions in the nNPC were selected consistent with those in the NE-NPC in spite of slight deviations in the map densities. Average diameters (means ± SD) are shown for each ring. **C** Plot of the diameter measurement. Dots indicate measurements on individual rings; the median and upper and lower quartile are indicated by black lines. **A-C** Sample: *S. cerevisiae* Not4Δ Nup188::GFP Nup170::mars.

**Table S1: *Drosophila* CRISPR information, related to „Generation of fluorescently tagged transgenic flies"**

| Fly line name | tagged protein (Chromosome location) | tagged terminus | linker | tag | guide RNA | injected fly line |
|---|---|---|---|---|---|---|
| Nup358::GFP | Nup358 (3R) | C | 4x(GGS) | mEmerald | 5'-GAACAAGAATGACGACGCGC-3' | *y[1] sc[*] v[1] sev[21]; P{y[+t7.7] v[+t1.8]=nanos-Cas9.R}attP40* (BDSC-78781) |
| Nup358::VN | Nup358 (3R) | C | 4x(GGS) | Venus N-terminus | 5'-GAACAAGAATGACGACGCGC-3' | *y[1] sc[*] v[1] sev[21]; P{y[+t7.7] v[+t1.8]=nanos-Cas9.R}attP40* (BDSC-78781) |
| Nup358::VC | Nup358 (3R) | C | 4x(GGS) | Venus C-terminus | 5'-GAACAAGAATGACGACGCGC-3' | *y[1] sc[*] v[1] sev[21]; P{y[+t7.7] v[+t1.8]=nanos-Cas9.R}attP40* (BDSC-78781) |
| Nup214::GFP | Nup214 (2R) | C | 4x(GGS) | mEmerald | 5'-gCAGGAATCGGGACTCTCTT-3' | *w[1118]; PBac{y[+mDint2] GFP[E.3xP3]=vas-Cas9}VK00027* (BDSC-51324) |
| mCherry::Nup358 | Nup358 (3R) | N | 3x(GGS) | mCherry | 5'-gTTTACAACGCGAAAAGAAG-3' | *yw; nos-Cas9(II-attp-40)* |
| GFP::Nup98 | Nup98 (3R) | N | - | mEmerald | 5'-GGCGGCGCGAAACCCAGCTT-3' | *yw; nos-Cas9(II-attp-40)* |
| mCherry::Nup153 | Nup153 (X) | N | 3x(GGS) | mCherry | 5'-gAGGTGAGTCTGAATCTAAAG-3' | *w[1118]; PBac{y[+mDint2] GFP[E.3xP3]=vas-Cas9}VK00027* (BDSC-51324) |
| GFP::Nup133 | Nup133 (3R) | N | - | mEmerald | 5'-GCAACAGCAAACAATCTAGC-3' | *y[1] sc[*] v[1] sev[21]; P{y[+t7.7] v[+t1.8]=nanos-Cas9.R}attP40* (BDSC-78781) |

### Table S2: *Drosophila* cryoET data acquisition and map parameters, related to Figure 3

| | | | | | | | |
|---|---|---|---|---|---|---|---|
| Microscope | Titan Krios G4, Selectris X energy filter, E-CFEG | | | | | | |
| Voltage | 300 kV | | | | | | |
| Camera | Thermo Scientific Falcon 4 | | | | | | |
| Nominal Magnfication | 53000 | | | | | | |
| Pixel size | 2.414 Å/px | | | | | | |
| Targeted total electron dose | 150 e-/Å^2 | | | | | | |
| Tilt-scheme | Dose symmetric around 11° pretilt, 2 ° steps, stage tilt range: -49 to +71 or +49 to -71 | | | | | | |
| Target dose rate | 5 e-/px/s | | | | | | |
| Targeted Defocus | Range: -1 to -5 µm | | | | | | |
| Energy Filter width | 10 eV | | | | | | |
| Acquisition software | SerialEM Version 4.0.14 and 4.0.20 | | | | | | |
| Alignment of tilt series | AreTomo Version 1.3.3 | | | | | | |
| Defocus estimation | gctf v1.06 | | | | | | |
| Tomogram reconstruction for TM | novaCTF | | | | | | |
| Tomograms used for TM | 13 | | | | | | |
| Tomogram reconstruction for STA | AreTomo Version 1.3.3 | | | | | | |
| NPC type | NE-NPC | | | | cNPC | | |
| Tomograms used for STA | 110 | | | | 10 | | |
| Initial # of NPC | 318 | | | | 34 | | |
| Map type | CR | IR | NR | NB | CR top* | IR | CR bottom** |
| Final # of particles | 1627 | 1629 | 1627 | 1626 | 202 | 202 | 202 |
| Resolution (Å) at FSC = 0.5 | 34.5 | 31.1 | 34.5 | 43.9 | 50.8 | 37.1 | 46.0 |

***** in CR position
** in NR position

**Table S3: Yeast cryoET data acquisition and map parameters, related to Figure 5**

| | | | | | | |
|---|---|---|---|---|---|---|
| Microscope | Titan Krios G2 with Gatan BioQuantum-K3 imaging filter | | | | | |
| Voltage | 300 kV | | | | | |
| Camera | Gatan K3 | | | | | |
| Nominal Magnfication | 42000 | | | | | |
| Pixel size | 2.176 Å/px | | | | | |
| Targeted total electron dose | 150 e-/Å^2 | | | | | |
| Tilt-scheme | Dose symmetric around 8° pretilt, 2 ° steps, stage tilt range: -52 to +66 or +52 to -66 | | | | | |
| Target dose rate | 12 e-/px/s | | | | | |
| Targeted Defocus | Range: -2 to -5 µm | | | | | |
| Energy Filter width | 20 eV | | | | | |
| Acquisition software | SerialEM Version 4.0.4 | | | | | |
| Alignment of tilt series | IMOD Version 4.11.5 and 4.10.51 | | | | | |
| Defocus estimation | gctf v1.06 | | | | | |
| Tomogram reconstruction for TM | novaCTF | | | | | |
| Tomograms used for TM | 28 | | | | | |
| Tomogram reconstruction for STA | novaCTF | | | | | |
| NPC type | NE-NPC | | | nNPC | | |
| Tomograms used for STA | 40 | | | 40 | | |
| Initial # of NPC | 281 | | | 278 | | |
| Map type | CR | IR | NR | NR top* | IR | NR bottom** |
| Final # of particles | 1489 | 1776 | 1489 | 1654 | 1654 | 1653 |
| Resolution (Å) at FSC = 0.5 | 44.0 | 35.5 | 46.4 | 39.9 | 36.8 | 38.3 |

***** in CR position
** in NR position

**Video S1: CCC scores from template matching with the *Drosophila* and human cytoplasmic ring template, related to Figure 4.**

Overlay of the tomogram shown in Figure 4 with scores volume resulting from template matching of cytoplasmic ring search templates. The scores volumes are shown in colour scale 'magma' on top of a grey-scale tomogram. The first half of the video shows template matching with the *Drosophila* cytoplasmic ring template from this study, the second half depicts scores from template matching with a human cytoplasmic ring search template (EMD-14325).
